# The pH Dependence of Niclosamide Solubility, Dissolution, and Morphology Motivates Potentially Universal Mucin-Penetrating Nasal and Throat Sprays for COVID19, its Contagious Variants, and Other Respiratory Viral Infections

**DOI:** 10.1101/2021.08.16.456531

**Authors:** David Needham

**Author notes:** Corresponding author: David Needham PhD, DSc, Department of Mechanical Engineering and Material Science, Duke University, Durham, NC, 27708, USA.

## Abstract

**Motivation:** With the coronavirus pandemic still raging, prophylactic nasal and early treatment throat sprays could help prevent infection and reduce viral load. Niclosamide has the potential to treat a broad range of viral infections if local bioavailability is optimized as mucin-penetrating solutions as opposed to more traditional microparticle-based sprays that cannot penetrate the mucin.

**Experimental:** pH-dependence of supernatant concentrations and dissolution rates of niclosamide were measured in buffered solutions by Nanodrop-UV/Vis-spectroscopy for niclosamide from different suppliers, as precipitated material, and as cosolvates. Data was compared to predictions from Henderson Hasselbalch and precipitation pH models. Optimal microscopy was used to observe the morphologies of precipitated and converted niclosamide.

**Results:** Supernatant-concentrations of niclosamide increased with increasing pH, from 1.77uM at pH 3.66 to 30uM at pH 8, and more rapidly from 90uM at pH8.5 to 300uM at pH 9.1, reaching 641uM at pH 9.5. Logarithmic rates for dissolution increased by ∼3x for pHs 8.62 to 9.44. However, when precipitated from supersaturated solution, niclosamide equilibrated to much lower final supernatant concentrations, reflective of more stable polymorphs at each pH that were also apparent for niclosamide from other suppliers and cosolvates.

**Conclusions:** Given niclosamide’s activity against COVID19, its more contagious variants, and other respiratory viral infections, these niclosamide solutions, that *put the virus in lockdown,* could represent universal prophylactic nasal and early treatment throat sprays. As solutions they would be the simplest and potentially most effective formulations from both an efficacy standpoint as well as manufacturing and distribution, with no cold chain. They now just need testing.

## 1. Introduction

This article lays out the case for new universal prophylactic nasal and early treatment throat sprays for COVID19, its more contagious variants (1), and other respiratory viral infections. The sprays are based on aqueous solutions of the anti-helminthic drug niclosamide, a drug with relatively low water solubility at pH 7 (and hence prohibitively low bioavailability). A preformulation drug characterization of niclosamide (pKa, water-solubility, and LogP) was carried out along with an experimental evaluation of the amount of niclosamide that could be dissolved in aqueous solution as a function of pH. Results revealed that the concentration of niclosamide in aqueous solution can be increased by simply increasing solution pH by only one to two logs, into the pH 8-9 range. However, studies also showed that, in the presence of solid phase niclosamide, the amount of niclosamide in solution is determined by the solid state nature of the material that the aqueous solution is in equilibrium with or is moving towards (2). And not every source of niclosamide is equal in polymorphic structure, solubility, and therefore *in vivo* therapeutic performance. Such solid phase material includes precipitated or particulate niclosamide as micronized or stabilized microparticles. Our lead niclosamide formulation is therefore a simple buffered mucin-penetrating solution with optimized bioavailability because it does not contain any microparticles of niclosamide. It represents a low dose prophylactic nasal spray that, according to recent literature on niclosamide and viruses, could stop virus replication at its point of entry in the nose, and a higher concentration throat spray, that could reduce viral load as it progresses down the back of the throat.

### 1.1. Niclosamide inhibits three of the six stages of viral infection

Why niclosamide? Niclosamide, marketed as Yomesan by Bayer (3) and other generics, has been routinely given to humans for the past 60 years as oral tablets in a 2-gram dose to cure cestode parasites (tape worms) (4). Over the past 10 years, it has emerged from multiple drug screens as a very interesting compound; not just as a pesticide but also with potential for cancer and many other diseases and conditions (5), and now a broad range of viral infections (6–21). The reason for this broad-range activity is that, because viruses are obligate intracellular pathogens, they cannot replicate without utilizing the machinery and metabolism of a host cell. As discussed by Goulding (22), there are six basic stages that are essential for viral replication at the cellular level: 1. *Attachment, 2. Penetration, 3. Uncoating, 4. Replication, 5. Assembly,* and *6. Release* of the mature virions. Niclosamide can inhibit three of these, namely, *<u>3. Uncoating</u>* that prevents RNA release from the endosome; *<u>4. Replication</u>*, where it reduces the amount of ATP available from mitochondria and so inhibits the viral transcription and translation events that are ATP dependent; and *<u>5. Assembly</u>* in the Golgi that then results in the secretion of non-competent virions.

While many current anti-virals either attempt to disrupt the synthesis and assembly of viral proteins or target host proteins and mechanisms required by the viral replication cycle (23), reformulated niclosamide offers a different, and potentially very effective, way to prevent and combat early viral infection. As discussed in more detail in a white paper in preparation (24), it enters cell membranes as a lipophilic anion where it acts as a proton shunt (11), dissipating pH gradients across a range of host cellular-organelle membranes, including mitochondria, endosomes and lysosomes, and even the Golgi. In the endosome it blocks entry of viral RNA (19) by not allowing the endosome pH to acidify, thereby preventing the acidic-pH-dependent conformational change required by the coronavirus spike protein to fuse with the endosome membrane (25, 26)). Similarly, as shown for other viruses it could potentially inhibit coronavirus assembly in the Golgi (27). One of its main effects though is in the inner mitochondrial membrane. Here it dissipates the pH gradient required to drive ATP synthase (11). This, in turn, reduces the cell’s production of its main energy molecule, adenosine triphosphate (ATP). Niclosamide’s oxidative phosphorylation effects are therefore enabled by a reduction in ATP that is upstream of many key cellular processes including transcription and translation of viral RNA (28). ATP is also a substrate for the multi-subunit enzyme RNA polymerase that adds ATP and the other ribonucleotides to a growing RNA strand. Furthermore, as described by Zimmerman et al, (29), ATP serves as a cofactor for signal transduction reactions using a variety of kinases as well as adenyl cyclase. Hence, the production and presence of ATP in every cell is essential to their functioning on multiple levels. Normally, cellular ATP concentration is maintained in the range of 1 to 10 mmol/L. However, titrating with niclosamide reduces the ATP content as readily measured using assays for cell “viability”, such as Cell Titer Glo (30). In *vaccinia poxvirus*, virus production actually requires increased amounts of ATP (31, 32), and so, if this holds for coronavirus, niclosamide could provide an even greater inhibitory effect.

Thus, niclosamide, at relatively low concentrations, in only the tens of micromolar range, basically *turns down the dimmer switch* on the cell’s energy production while still preserving reversible viability (33). As such, niclosamide can inhibit virus replication as demonstrated in Vero 6 cells at only 1µM (10) and has also been shown to have similar anti-replicative efficacy in cultured Calu-3 lung cells, at 2µM (34). Data is still needed on actual airway epithelial cells which is essential to show for its potential use in COVID prevention and early treatment.

### 1.2. The Need for New Formulations

Because of these quite dramatic and potentially efficacious effects, and the urgency of the COVID19 pandemic, in addition to simply repurposing the original oral tablets, there have been efforts to develop new formulations and routes of administration for niclosamide, to validate them in preclinical studies, and clinically test niclosamide for COVID-19 (see this reference to clincaltrials.gov (35)). These trials include new intramuscular injections by Daewoong Pharmaceuticals of South Korea (36), and a nasal spray and inhalant as a spray-dried-lysozyme particulate of micronized-niclosamide (37), that has entered a 1,500 high-risk kidney patient study in the UK with Union Therapeutics in Denmark (38).

However, regarding the nasal spray, microparticles are not expected to penetrate the mucin barrier that protects the respiratory epithelial cells (39–41), and so this now motivates a deeper physicochemical evaluation of niclosamide and especially how to increase its concentration in solution using simple buffers to access the epithelial cells. The goal of this study is therefore to optimize the bioavailability of niclosamide as a mucin-penetrating solution and saturate the first layer of epithelial cells that are the targets for SARS-CoV-2 viral infection (42).

#### 1.2.1 The main issue is optimization of drug bioavailability

The main issue for any formulation of niclosamide (and indeed other BCS Class II and IV low water-solubility drugs) is how to optimize local bioavailability. This is especially important in the nose and throat. There are two parts to this: the intrinsic low solubility of niclosamide in aqueous solution compared to its efficacy; and the fact that the more traditional choice for nasal sprays of microparticles cannot penetrate the mucin (39–41).

While its efficacy at inhibiting viral replication has been measured in Calu-3 cells to be 2uM - 3uM (6, 34), the amount of niclosamide that can be dissolved in aqueous solution at neutral or lower pH, as encountered in the nasal pharynx (43), is of the same order, or perhaps slightly less, i.e., ∼1uM - 2uM, ((2) and as measured here). Because of this low aqueous solubility, one might imagine that intranasal microparticle delivery would increase the amount delivered. However, the mucosa is designed to protect the nasal epithelial cells and prevent the permeation of particles with a size greater than a few 100 nanometers (41). As a result, any microparticle formulation cannot deliver molecular drug directly to the epithelial cells where initial infection takes place because of the protective mucin barrier that is produced by, and covers, these cells (41). This means that any drug microparticle suspension can only provide the fraction of drug that is soluble in aqueous solution in any delivered suspension. This may still show efficacy depending on the solubility-to-efficacy ratio. A good example is Flonase (44) and Nasonex (45, 46) (see also ***Supplemental Information S1.2***). Here, solubilities of their APIs, fluticasone propionate and mometasone furoate, are 21uM and 23uM, respectively. By comparison, the relative trans-activation potency (EC_50_) for fluticasone propionate is 0.320 ± 0.04 nmol/L and that for mometasone furoate is even less at 0.069 ± 0.021 nmol/L. That is, the amount of drug required to induce efficacy is 6,000 and 30,000 times greater in bioavailable aqueous concentration of the drugs compared to their nanomolar efficacy. However, for the low solubility niclosamide, this solubility-to-efficacy ratio is 1 or less (***see also Supplemental Information S1.3***). Microparticle niclosamide formulations may show some efficacy but are therefore clearly not optimal. There is therefore a need for a more bioavailable solution.

#### 1.2.2 Meeting the challenge with a simple solution

It is here that we have focused on what it takes to provide a mucus-penetrating niclosamide solution rather than microparticulate material that is simply trapped at the mucus surface, and then excreted as the mucin is replaced (every 15-20 minutes (47)). We have found a way to increase the solubility of niclosamide and therefore its bioavailability to infectable and infected epithelial cells just by slightly increasing the pH in simple aqueous buffered solutions (48). This now optimizes the delivery of niclosamide in a low dose prophylactic nasal spray and as a higher dose early treatment throat spray.

As described in Results and Discussion, at pH 7.9, a 20uM niclosamide nasal solution is within the natural pH range for the nasopharynx and, at pH 9.1, a higher concentration 300µM throat spray is at a tolerable pH for the oropharynx. These niclosamide solution concentrations are ∼10x to ∼150x above the *in vitro* inhibition of SARS-CoV-2 viral replication as measured to date in Vero 6 and Calu-3 cells of 1µM −3µM (6, 34, 49). Also, the preliminary studies by Kim (33) in airway epithelial cells, show that at an IC_50_ of 20uM to 30uM niclosamide gives the necessary reductions in cell ATP, and that the lowered cell viability is reversible and does not result in any cell death. While the vaccine is highly protective against hospitalization and severe disease, break through infections are now occurring, and virus is being spread by the asymptomatic-vaccinated requiring new mask mandates, that all seem to be linked to the delta variant (50, 51). Prophylactic nasal and early treatment throat sprays could *“put the virus in lockdown”* or not even let the viral RNA into the nasal epithelial cells. As such, they could be a huge boost in the control of viral load, and reduce development of infection for unvaccinated and vaccinated spreaders.

The envisioned final products are solutions of niclosamide made up in pH buffer and available in 10mL doses in 15 mL nasal and throat spray bottles. The low dose 20uM niclosamide solution at pH 7.9 can be sprayed intranasally and the higher dose 300uM sprayed to the back of the throat.

##### Bottom line

With the coronavirus pandemic still raging, more contagious (delta and delta+) variants already in circulation (1), a world-wide average of only 15.4% fully vaccinated (52), and may parts of the world with <1% vaccinated (53) there is still an urgent need for additional mitigation efforts to support current and potential vaccinations. For unvaccinated (including children and now babies where COVID cases are now rising (54)), anti-mask, and anti-vaccination populations, an effective nasal prophylactic and early treatment throat spray would likely enhance those efforts and could be more compliant than mask wearing. Such simple sprays could also potentially be further *protection behind the mask* and even instead of the mask, if and when tested. Thus, while vaccines and antibody treatments are certainly effective, they are designed to work only after infection has taken place, i.e., they do not prevent infection, as we are now seeing with breakthrough infections even for vaccinated persons (50, 51). They are also limited to a particular virus, a particular viral protein, and may eventually need to be boosted or redesigned for an emerging viral variant. In contrast, Niclosamide is a universal host cell modulator and so is specific for every virally infected or infectable cell. As postulated by Laise et al (23), *“Targeting host cell mechanisms may have more universal and longer-term value because the same host factors may be required by multiple, potentially unrelated viral species and because host target proteins mutate far less rapidly than viral proteins, thereby limiting emergence of drug resistance”.* We concur; and would add that lipid bilayer membranes mutate even less.

If it can be reformulated and tested as a nasal and throat spray with optimized bioavailability, niclosamide could potentially represent a universal prophylactic and early treatment for many viral infections, including ones that mutate and escape current vaccine protection. When formulated as an optimized mucin-penetrating solution, niclosamide can potentially help to control initial infection and viral loads across the board, while vaccines and other oral antivirals provide subsequent immune protection. As with vaccine development, there is a need for billions of doses, (especially on a frequent basis) and so there is certainly room for several prophylactic formulations, tested and produced by multiple governments, infectious disease institutes, schools of public health, and companies that can help to control such initial viral infection and early symptoms. A low dose (20uM) prophylactic solution of niclosamide at a nasally safe and acceptable pH of 7.9 and a (up to 300uM) throat spray at pH 9.1 may be one of the simplest and potentially most effective formulations from both an efficacy standpoint as well as manufacturing and distribution, with no cold chain. This niclosamide solution formulation has already been scaled up to multi-liter volumes as a 503b pharmacy batch that can be provided in 10mL sealed and capped sterile vials; it just needs testing. The physico-chemical basis for this formulation and how to make it are now presented. “Why Niclosamide?”, and its intracellular *virostatic* effects, along with how these measurements are translated into spray products and preclinical and clinical outcomes are discussed in more detail in a white paper in preparation (24).

## 2. Research Design

We have much previous experience working with and preparing niclosamide for drug delivery, including as a prodrug therapeutic for anti-cancer applications (55, 56). Following publication of the bioRxiv preprint by Jeon et al in March 2020 (10) (now peer-reviewed (49)), that demonstrated anti-viral activity of niclosamide against SARS-CoV-2, our efforts were refocused on ways to increase the amount of niclosamide in sprayable nasal and throat suspensions (48). A description of the initial thinking that went into the work and that led to the solution formulation is given in the ***Supplemental Information S1***. It is an interesting account that, along with this Research Design section, should help to inform, especially students and post docs, what went into the research and development. It includes: more on the case of Nansonex and Flonase; that Niclosamide’s solubility at physiologic pH is actually in the same range as its efficacy; could Niclosamide be encapsulated or stabilized as nanoparticles?; and given the “solution” to the problem, how much niclosamide is actually sprayed per dose?

In preparation for designing and making any drug delivery system, it is also important to carry out a preformulation drug characterization (see ***Supplemental Information S2***). Such analyses are focused on the basic molecular properties of the niclosamide, including its pKa, the solubility of its protonated (S_o_) and deprotonated (S_-ve_) forms, the resulting pH-dependent amount of niclosamide in solution, and its octanol water partition coefficients (LogP, and the pH-dependent Log D) that underly the ability of niclosamide to partition into the various lipid bilayer membranes and exert its proton shunt effect.

### 2.1 Overall Goal and Aims

The overall scientific goal of this study was to explore the expected pH-dependence of Niclosamide implied by the literature and calculation databases for its pKa (11, 57–59). This indicated that the amount of niclosamide in aqueous solution should increase with increasing pH. Experiments were therefore carried out to evaluate this pH dependence and compare it to theory for a sourced niclosamide from a chemical supplier, (AK Sci, CA). The initial aims therefore were to measure the equilibrium pH dependence of niclosamide (AK Sci) in a series of buffered solutions (nominally pHs 4 – 9.5) and its rate of dissolution in the same buffer series. However, on occasion, supernatant solutions that had attained equilibrium with the slight excess of solid material, showed a reduction in concentration, indicating that some samples were moving towards a lower solubility solid form. A second series of experiments were therefore deigned to explore these conversions that are already known for niclosamide (2, 60–62) but have not been investigated as a function of pH. Equilibrium supernatant concentrations and dissolution rates for niclosamide from a second supplier, (Sigma), were therefore investigated, as were precipitated samples and specially prepared acetone and ethanol cosolvates. As reported in Results, while the Sigma sample initially produced similar dissolution and supernatant concentrations as the AKSci material, when left overnight, it converted to a lower solubility polymorph. Similarly, when precipitated at each pH, there was still a pH dependence to the amount in solution but now at a much lower level than the as-supplied AK Sci niclosamide powder. The cosolvates also provided a lower amount of niclosamide in supernatant solution.

### 2.2 Specific Aims

Experimentation therefore focused on five main specific aims (SAs).

#### SA 1. To measure the pH dependence of AK Sci Niclosamide in buffered solutions

As predicted by Henderson Hasselbalch and precipitation pH models, the first task was to confirm that the amount of niclosamide in buffered solution increased with increasing pH over the nominal pH range 4 to 9.5. Small excesses of powdered niclosamide from a commercial supplier (AK Sci, CA) were equilibrated in pH buffers in stirred vials and the filtered supernatant niclosamide concentration was measured using a calibrated nanodrop UV/Vis protocol.

#### SA 2. To measure dissolution rates of AK Sci niclosamide as a function of pH including a new methodology for measuring the supernatant concentration in situ

In a series of dissolution measurements, a new methodology was established for measuring the supernatant concentration of niclosamide *in situ* using UV/Vis absorption of stirred niclosamide solutions equilibrated with supplied powdered niclosamide. This new sampling technique utilized a nanodrop UV/Vis spectrophotometer for detection of niclosamide at 333nm and 377nm using only 2uL samples drawn directly from the stirred suspensions. It allowed the measurement of initial dissolution rates and full dissolution curves *in situ* as the drug was dissolving versus time. Dissolution time periods were 60 minutes and longer for a series of pH solutions using the AK Sci niclosamide, an optimized mixing protocol was also developed.

#### SA 3. To measure equilibrium supernatant concentrations and dissolution rates of Sigma niclosamide as a function of pH

Similar experiments were carried out for the Sigma niclosamide as above. Excess 1mM nominal concentrations were equilibrated with pH 9.5 buffer using the optimized stirring protocol over a period of 3 hrs and equilibrium supernatant concentrations were measured by nanodrop UV/Vis spectroscopy. A series of samples were made over the pH range 7 to 9.5 and allowed to continue stirring overnight. The new equilibrium concentrations were measured after filtering through a 0.22um filter. Optical microscope images were obtained of particle morphologies at each pH.

#### SA4. To measure equilibrium supernatant concentrations when Niclosamide was precipitated at the same pHs

When precipitated from supersaturated solution the formed microcrystals are expected to take on their “natural” morphology associated with nucleation and growth at that pH. Using the solvent injection technique, it was possible to obtain such precipitated material and measure their equilibrium niclosamide solution concentrations corresponding to precipitation at each pH. Optical microscopy again allowed images to be obtained of the particle morphologies at each pH.

#### SA 5. To evaluate dissolution rates and equilibrium niclosamide concentrations for niclosamide- “cosolvates” (water, ethanol and acetone)

The above studies led to a series of experiments that sought to provide additional information as to the nature of the commercially available niclosamide material that dissolved so readily (AK Sci and Sigma), but where, in one case (Sigma) it converted to a lower solubility form while in the other (AK Sci) was mostly stable at high concentration for days in equilibrium with its original powder. Niclosamide was therefore precipitated into water to obtain the fully hydrated form or was recrystallized from ethanol and acetone and then dissolved to equilibrium in pH 9.3 buffer to determine their dissolution profiles, final niclosamide concentrations, and particle morphology, again by optical microscopy. Note: a similar protocol was used to make them as reported by van Tonder et al, (62). Subsequent characterization by (Burley et al Nottingham (63)) using Raman spectroscopy/microscopy, infra-red spectroscopy, X-ray diffraction (powder and single crystal), thermal methods, will show if they are truly cosolvates.

## 3. Experimental

### 3.1 Materials

Niclosamide was from AK Sci (CA) (Lot No. 90402H, listed as, at least 98% pure by HPLC) and from Sigma, (N3510); Water was deionized and filtered through 0.22um filters. TRIZMA buffer was from Supelco and comprised: Trizma Base, 99.8+%, reagent grade Tris(hydroxymethyl)-aminomethane(HOCH_2_)_3_CNH_2_, Mol Wt 121.14g/mol, white crystalline powder; Trizma HCl, 99+%, reagent grade (Tris[hydroxymethyl] aminomethane hydrochloride), (HOCH_2_)_3_CNH_2_•HCI Mol Wt 157.60g/mol, white crystalline powder. Ethanol KOPTEC 200% proof and acetone BDH 1101 were from VWR. pH buffers were made using the TRIZMA HCl and TRIZMA base buffer system (Supelco). From a 1L stock solution at pH 9, a series of 100mL buffer solutions in 250mL glass screw capped bottles were made nominally at pHs of 4.0, 7.0, 7.5, 8.0, 8.25, 8.5, 8.75, 9.0, 9.3 and 9.5 At the highest TRIZMA base-concentrations it was possible to exceed pH 9 and extend the range to pH 9.5. The pH solutions were made by adding to the stirred pH 9.5 solution suitably small amounts of 2M HCl using a pipettor and fine-tuned with 0.1M to achieve the lower pHs or by adding small amounts of Trizma base to re-raise the pH. pHs were measured using a Mettler SevenEasy™ pH Meter S20, calibrated prior to any measurements using standard (VWR) buffers of pH 4, 7 and 10. Nominal pH 4 buffer was made using sodium citrate-citric acid buffer.

AKSci niclosamide purity was measured by HPLC-UV Chromatogram for Niclosamide in MeOH and was found to be 98.4% pure based on %-area, and so agreed with the AKSci data.

Niclosamide was also found to be stable in a Forced Degradation test, where solution of niclosamide was prepared at 100 µg/mL in 1 N NaOH, stored at room temperature, and periodically analyzed by LCMS over a 5-day period. Degradation to its breakdown products of 2-chloro-4-nitroaniline and 5-chloro-2-hydroxybenzoic acid were observed at this high pH. However, at neutral pH and pH 9, the solutions were stable as niclosamide for at least 16 days of measurement.

### 3.2. Methods

#### 3.2.1 Solvent exchange technique for making calibration standards and supersaturated solutions

The solvent exchange technique, normally used to make nanoparticles (55), can readily be used to make small volumes of niclosamide solutions. The procedure is to simply inject a relatively concentrated ethanolic niclosamide solution into an excess of anti-solvent, i.e., the stirred buffer. ***Figure 1*** shows the eVol syringe, 20mL scintillation vial, and magnetic stirrer set up.

**Figure 1.**
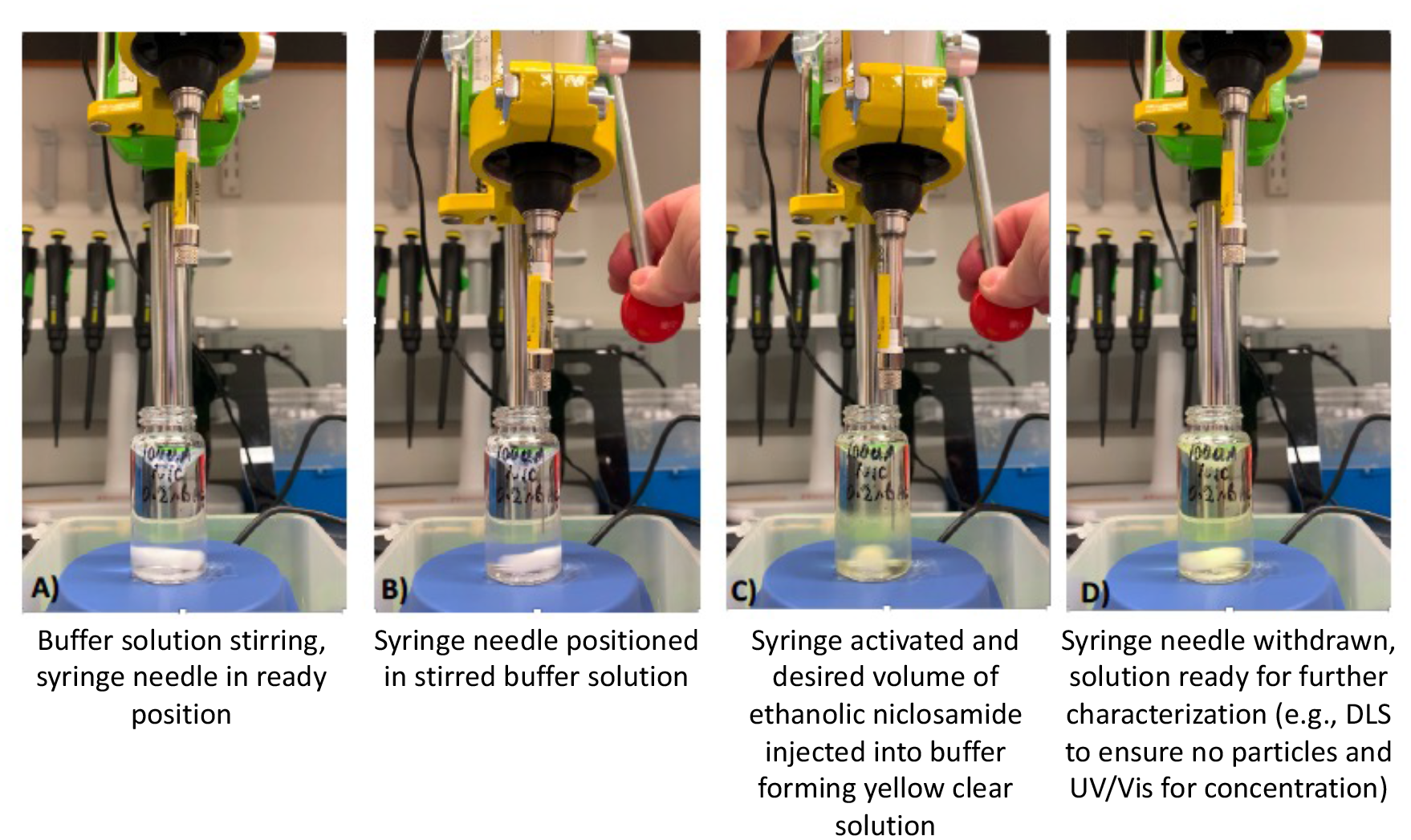
eVol syringe mounted in drill press, with 20 mL scintillation vial and magnetic stirrerset up, showing the simple procedure for introducing microliters of ethanolic niclosamide into the stirred vial.

Basically, the technique involves diluting a concentrated ethanolic niclosamide solution and exchanging the good solvent (ethanol) for the anti-solvent (aqueous buffer) but doing it in such a way that the final concentration does not exceed the solubility limit of niclosamide in the buffer at that pH. As a result, there is no precipitation of niclosamide (this was checked via Dynamic Light Scattering, (DLS)). As shown in ***Figure 1***, an eVol syringe (Trajan Scientific and Medical. Trajan Scientific Australia Pty Ltd) is clamped securely in a commercially available drill press (Yeezugo, Guangzhou, China). This mounting allows for repeatable and accurate positioning of the syringe needle tip in the stirred solution, --a feature that is essential for injection of the solution into the most effectively stirred volume of the solution in the vial. While less of an issue when making solutions, repeatable and consistent injection speed and mixing environment is critical for supersaturation precipitation since the vortex is not the most efficiently stirred part of the system (64) and particle nucleation is very sensitive to the mixing environment (55). The protocol for making the standards or precipitating material is as follows:

Withdraw 12 milliliters of the buffer solution into a 30 mL BD syringe. Insert and luer–lock a 0.22 um filter to the syringe and prime the filter by expelling ∼2 mLs of solution into a waste beaker. Add 9.9 mLs of the solution from the syringe through the filter into the 20 mL scintillation vial. Load the eVol syringe with 100uL of a desired niclosamide solution in ethanol that has also been filtered through a 0.22um filter to remove any insoluble residual particulate material in the supplied niclosamide powder.

Add an ethanol-cleaned, small magnetic stir bar to the vial and place the vial on a magnetic stirrer as shown in the photographic images in ***Figure 1***. Turn on the magnetic stirrer to create a small vortex (***Figure 1A***). Bring down the syringe needle and position it over to the side of the vortex (***Figure 1B***) i.e., for better mixing the syringe tip needs to be close to the ends of the stir bar. For the stock standard, inject the 100uL of 30mM ethanolic Niclosamide into the 9.9mLs of stirred pH 9.3 buffered solution at moderate injection rate (***Figure 1C***). Within 1 to 2 seconds of finishing the injection, turn off the stir motor, raise the needle (***Figure 1D***) and remove the vial. This method makes 10mLs of 300uM Niclosamide including 1% residual ethanol. For precipitation of niclosamide from supersaturated solution the desired concentration of ethanolic niclosamide was used to provide a final niclosamide concentration that was above the solubility of niclosamide in solution at a particular pH, as described below in 3.2.4. For more details on calibration standards including results, see ***Supplemental Information S3***.

#### 3.2.2 Equilibrium Dissolution of AK Sci Niclosamide versus pH

A series of equilibrated niclosamide solutions were prepared at each nominal pH from 4 to 9.5 (4, 7, 7.5, 8, 8.25, 8.5, 8.75, 9, 9.3, 9.5) by dissolving the AK Sci niclosamide at concentrations that were in excess of its expected solubility, i.e., 100uM (0.33mg/10mLs) for pH 4, 7, 7.5 and 8; 300uM (1mg/10mLs) for pH 8.25, 8.5 and 8.75) and 1mM (3.3mg/10mLs) for pH 9, 9.3 and 9.5. As can be appreciated, again, very small amounts were required, (∼0.33mg - 3.3mgs). Niclosamide, as obtained directly from the supplier (AK Sci) was weighed into a weigh boat and added to 10 mLs of each buffer solution in 20mL screw top scintillation vials.

The solutions and undissolved particles in the vials where then briefly shaken by hand to wet and immerse the hydrophobic niclosamide powder and stirred by magnetic stirrers. UV-Vis measurements were made a few hours after making and stirring, and after 1-8 days to ensure equilibrium. Absorbance was compared to a standard calibration curve (given in ***Supplemental Information S3***). It is important to show this because the pH 9.3 spectra are actually new.

Since some suspensions were visibly cloudy due to the excess undissolved material, in order to avoid sampling particles that could interfere with the UV/Vis measurement, 0.5mLs of the supernatants were taken, spun down by Eppendorf-centrifugation (10mins at 22,000G) to remove any suspended particles, and analyzed in 2uL samples by UV/Vis nanodrop spectrophotometer (ThermoFisher 1000). Full spectra were recorded and the absorbance at the 333nm peak was compared to the calibration in order to determine the niclosamide concentrations of the supernatant. Their final pH was remeasured on a Mettler Toledo pH meter.

For the UV/Vis measurements, each buffer was used as its own blank, and at least 5 individual samples were taken for each buffer and averaged to establish the “blank baseline” which was usually between 0.000 and 0003 absorption values. These values were subtracted from the niclosamide measurements. This subtraction was particularly important for the very low niclosamide concentrations at pHs 3.66 and 7, where the signal was very close to, but still distinguishable from, the usual noise of the instrument.

#### 3.2.3 Rates of Dissolution for AK Sci Niclosamide powder

The dissolution of niclosamide powder was measured in a timed dissolution study using the nanodrop spectrophotometer. In order to try and standardized the particle size and therefore surface area of the niclosamide particles, the as-received powder was ground with a mortar and pestle. This was actually done for all samples, (from Sigma, the water-precipitated niclosamide, and the ethanol and acetone cosolvates). Appropriate amounts of AK Sci niclosamide powder, as in the above equilibration studies were weighed into dry 1.5mL Eppendorf tubes and capped. Since, as shown in results, final concentrations of niclosamide for pH 7 to 8.25 were still relatively low, only pH 8.5, 8.75, 9, 9.3 and 9.5 were tested. 10mLs of each buffer were aliquoted from a 10mL BD syringe fitted with a 0.22um filter into 20mL scintillation vials, and a cleaned magnetic stirbar was added to the vial. At time (t) = 0, the contents of one of the Eppendorf’s was rapidly emptied into the buffered solution vial. The vial and solution were quickly shaken by hand in order to wet and immerse the hydrophobic powder into suspension, and the vial was placed on the magnetic stirrer. 2uL samples for UV/Vis measurement on the nanodrop spectrophotometer were taken by a 2-20uL pipettor at regular time intervals, e.g., every 30s up to 7 minutes, every minute from 8 - 16 mins, every 2 mins from 18-30 mins and then every 5 mins from 30 – 60 mins. Thereafter, samples were taken at 90, 120, and 180 mins, as well as the next day.

Here, a new technique for measuring drug dissolution was developed. Rather than having to employ large volumes of suspension and a series of tubes, connectors, pumps, filters, and stirred cuvettes, 2uL samples were taken directly from the 10mLs of suspension in a stirred 20mL vial. This allowed direct *in situ* measurements to be made of the dissolution of the various niclosamide samples as a function of time at each pH (8.5 – 9.5). This gave their initial rates of dissolution including more optimized mixing conditions for the supplied material. By utilizing 2uL sampling, on a limited excess of powdered material in the stirred supernatant and making measurements onthe nanodrop spectrophotometer, it was very rare for the 2uL volume to include a powder particle. In any event, if this happened, the nanodrop software identified it as perhaps a bubble or error due to changed path length, and the measurement could be excluded and repeated.

#### 3.2.4 Supersaturated Solution Precipitation of Niclosamide: Solubilities at Each pH and Corresponding “Natural” Morphologies

This next experiment was carried out in order to determine the amounts of niclosamide in solution corresponding to what are expected to be the “natural” morphologies of niclosamide when precipitated at each pH. Using a series of stock solutions in ethanol, final supersaturated concentrations of niclosamide were created in each pH buffer in slightly excess amounts, i.e., just enough to generate precipitated material given that time was required for the stochastic nucleation and precipitation to occur. This excess, kinetically soluble, niclosamide was mixed into the buffers using the solvent injection technique. 33uL of 30mM into 10 mls of buffer gave supersaturated solutions of 100uM for pH 3.66, 7, 7.5, 8, 8.25; 100uL of 30mM gave a 500uM supersaturated solution for pH 8.5 and 8.75; and 333uL of 30mM gave a 1mM supersaturated solution for pH 9.0, 9.3 and 9.5. The solutions were again stirred until precipitation was observed, often overnight and sometimes longer. In the case where no precipitate was seen in the pH 9.5 solution after 1-2 days stirring, additional volumes of 30mM ethanolic niclosamide were added and the solution continued stirring until a precipitate was obtained. For this pH 9.5 sample, solution concentrations had to be increased to 3mM supersaturation and even then, the kinetic solution was stable for at least a few hrs at 3mM niclosamide. The precipitated suspensions were examined by optical microscopy using bright field optics, Köhler illumination, and a 40x objective to obtain micrograph images of the precipitates. 0.5mLs of supernatants were centrifuged in 1.5ml Eppendorf tubes for 10 mins at 22000 G to obtain clear supernatant solutions that were in equilibrium with the various precipitated morphologies. UV/Vis spectra and 333nm intensities were again obtained by the nanodrop spectrophotometer.

#### 3.2.5 Dissolution of Niclosamide Solvates

To start to evaluate the various expected polymorphs and to try and obtain a morphology like the purchased materials (of unknown processing), three samples were made i.e., the “water-hydrate” precipitated from ethanol into excess water, and two recrystallized samples from acetone and ethanol presumed to be “cosolvates” because the method of making followed that by van Tonder et al (2, 60, 62) for their various cosolvates that were evaluated by x-ray diffraction, IR spectra, and calorimetry. These were made and evaluated as follows.

##### Water precipitate

Niclosamide was precipitated from supersaturated solution by solvent exchange into excess deionized water by injecting 400uL of ethanolic 25mM niclosamide solution into 10mLs of stirred pH 9.3 buffer, (3.8% residual ethanol) i.e., much as might be done after synthesizing the niclosamide dissolving in an ethanol solvent and recovering by precipitation into excess water. The niclosamide precipitated immediately as its usual, visible by eye, white precipitate (see image in ***Figure S1*** Supplemental Information). It was filtered on a sintered glass filter, washed three times with deionized water, and dried. The material was ground by mortar and pestle ready for the dissolution test and viewed microscopically to evaluate its crystal morphology.

##### Niclosamide Recrystallized from Acetone and Ethanol

Niclosamide was recrystallized from acetone and ethanol by dissolving excess niclosamide into the solvent, gently warming in a fume hood to dissolve excess niclosamide, warmed to evaporate-off ∼50% of the solvent, that were then allowed to cool under stirring. The recrystallized niclosamide solvates were filtered and dried. Each material was also viewed microscopically to evaluate their crystal morphologies.

##### Dissolution of the solvates

As with the AK Sci and Sigma materials, ∼3.3mg of niclosamide from each of the hydrate and cosolvates was weighed and added to stirred pH 9.3 buffer to give a nominal 1mM niclosamide in suspension ready to dissolve. 2uL aliquots were taken directly from the stirred solution in the 20mL scintillation vial at appropriate time intervals (as above) and absorbance at 333nm was measured by nanodrop spectrophotometer.

#### 3.2.6 Dissolution of Sigma Niclosamide and its conversion to the low solubility form

A final experiment evaluated the dissolution of niclosamide from a second supplier, (Sigma), over the pH range 7 to 9.5. This powder was much lighter in color, a creamy yellow, as opposed to the browner yellow of the AK Sci product. As before (3.2.3) excess niclosamide was weighed to give an equivalent of 100uM (0.33mg) for pHs 7, 7.5 and 8.0; 200uM (0.65mg) for pHs 8.25 and 8.5, 500uM (1.65mg) for pHs 8.5 and 9.0 and 1mM (3.3mg) for pHs 9.3 and 9.5. When ready, each of these samples of powdered Sigma niclosamide was added to individual 10mLs of buffer solution in 20mL vials. As above, the vial and solution were quickly shaken by hand in order to wet and immerse the hydrophobic powder into suspension, and the vial was placed on the magnetic stirrer. The suspensions were stirred by magnetic stirbar for three hours. During this time, 2uL samples for UV/Vis measurement on the nanodrop spectrophotometer were taken by a 2-20uL pipettor at regular time intervals, e.g., every 30s up to 7 minutes, every minute from 8 −16 mins, every 2 mins from 18-30 mins and then every 5 mins from 30 – 60 mins.

At least 5 samples were taken to give the average niclosamide absorbance. Each pH buffer was used to blank the measurements and the blank values were again subtracted from the niclosamide absorbance values. Samples were left stirring overnight. At this point 0.5mls of each were centrifuged at 22,000G for 10 minutes and the supernatant niclosamide concentration was measured by UV/Vis nanodrop spectrophotometer.

## 4. Results

### 4.1 Equilibrium Dissolution of Niclosamide versus pH

20mL scintillation vials containing the equilibrated niclosamide solutions measured by UV/Vis at 333nm peak (by Thermo Scientific 1000 Nanodrop Spectrophotometer) as equilibrated stirred samples are shown in ***Figure 2***. It is clear, as with the calibration standards (see ***Supplemental Information S3.2***) that the characteristic yellow coloration of the solution gradually increased with increasing pH. A 300uM sample “standard” made by ethanol injection is shown for comparison (far right). In the final formulation, the preferred method of preparing the niclosamide-in-buffer solutions would be by ethanol injection of an ethanolic niclosamide solution into the buffer. This would allow the ethanolic solution to be pre-filtered to remove the small amount (<2%) insoluble material in the AKSci material.

**Figure 2.**
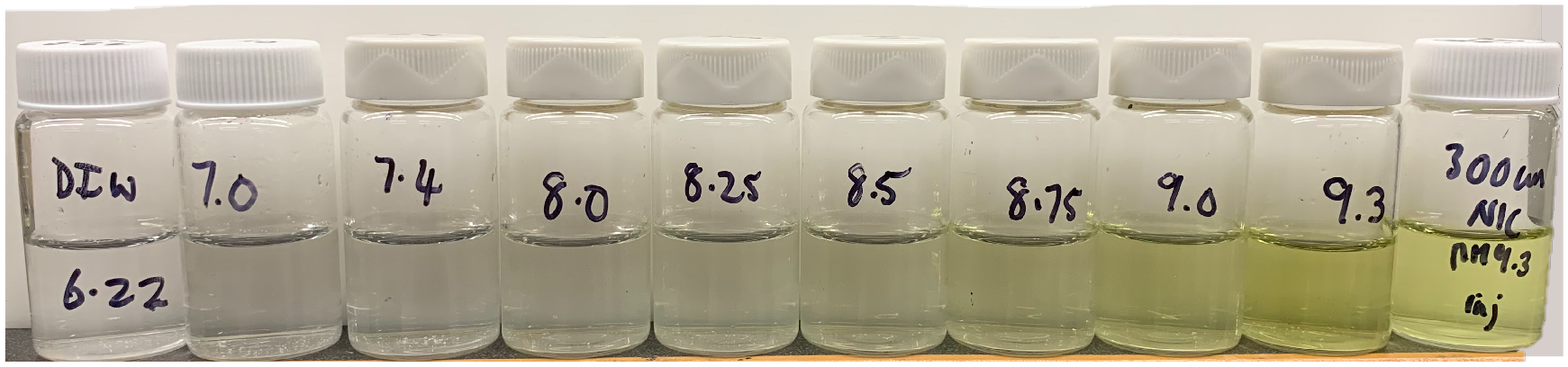
10mL samples of excess niclosamide dissolved in pH buffers (nominally 7.0 – 9.3). Also shown is a deionized water sample nominally pH 6.22 (far left,) and a 300uM Niclosamide in a pH 9.3 standard (far right), made by solvent injection (measured at 301uM ± 5uM). The image shows the increasing “yellowness” characteristic of niclosamide in solution. Excess powdered AK Sci niclosamide is seen at the bottom of each vial and, when stirred into the supernatant, was such a small amount that it was rarely included in the 2uL sampling.

For convenience, shown in ***Table I*** are the nominal and measured pH and average supernatant Niclosamide concentrations [Nic] (μM) corresponding to each of the vials in ***Figure 2***. Averages and Standard Deviations were taken from five UV/Vis measurements at each pH and the blanks subtracted. Additional re-checks to the original series were taken in the steep part of the curve at pHs 9.1, 9.3 and 9.5. After addition and equilibration of niclosamide in solution, the pH values were fairly stable compared to the nominal pH.

**Table I.** Supernatant concentrations of AK Sci Niclosamide (uM) measured by UV/VIS (nanodrop). Averaged are five measurements for each supernatant solution pH (nominal) and measured, for multiple samples after stirring to equilibrium (2-5 days), including re-checks to original series at pHs 9.1, 9.3, 9.5.

| (Nominal) and Measured pH | Average [Nic] (uM) (UV/Vis, 333nm) | Std Dev [Nic] (uM) |
| --- | --- | --- |
| (3.6) 3.66 | 1.77 | ± 1.0 |
| (7.0) 6.89 | 3.8 | ± 1.0 |
| (7.5) 7.4 | 7.4 | ± 1.0 |
| (8.0) 7.94 | 18.3 | ± 9.6 |
| (8.0) 8.08 | 19.5 | ± 12.4 |
| (8.25) 8.31 | 61.6 | ± 2.6 |
| (8.5) 8.47 | 87.0 | ± 1.3 |
| (8.75) 8.68 | 122.1 | ± 2.1 |
| (9.0) 8.92 | 228.9 | ± 5.7 |
| (9.1) 9.1 | 316.0 | ± 1.3 |
| (9.3) 9.24 | 414.9 | ± 2.6 |
| (9.3) 9.24 | 521.1 | ± 2.5 |
| (9.3) 9.31 | 446.4 | ± 2.1 |
| (9.3) 9.37 | 527.7 | ± 1.3 |
| (9.5) 9.31 | 581.6 | ± 8.1 |
| (9.5) 9.49 | 640.7 | ± 3.8 |
| (9.5) 9.56 | 640.8 | ± 4.2 |

The concentrations of these same supernatant solutions are plotted in ***Figure 3A*** for each sample (standard deviations are within the size of the symbols).

**Figure 3.**
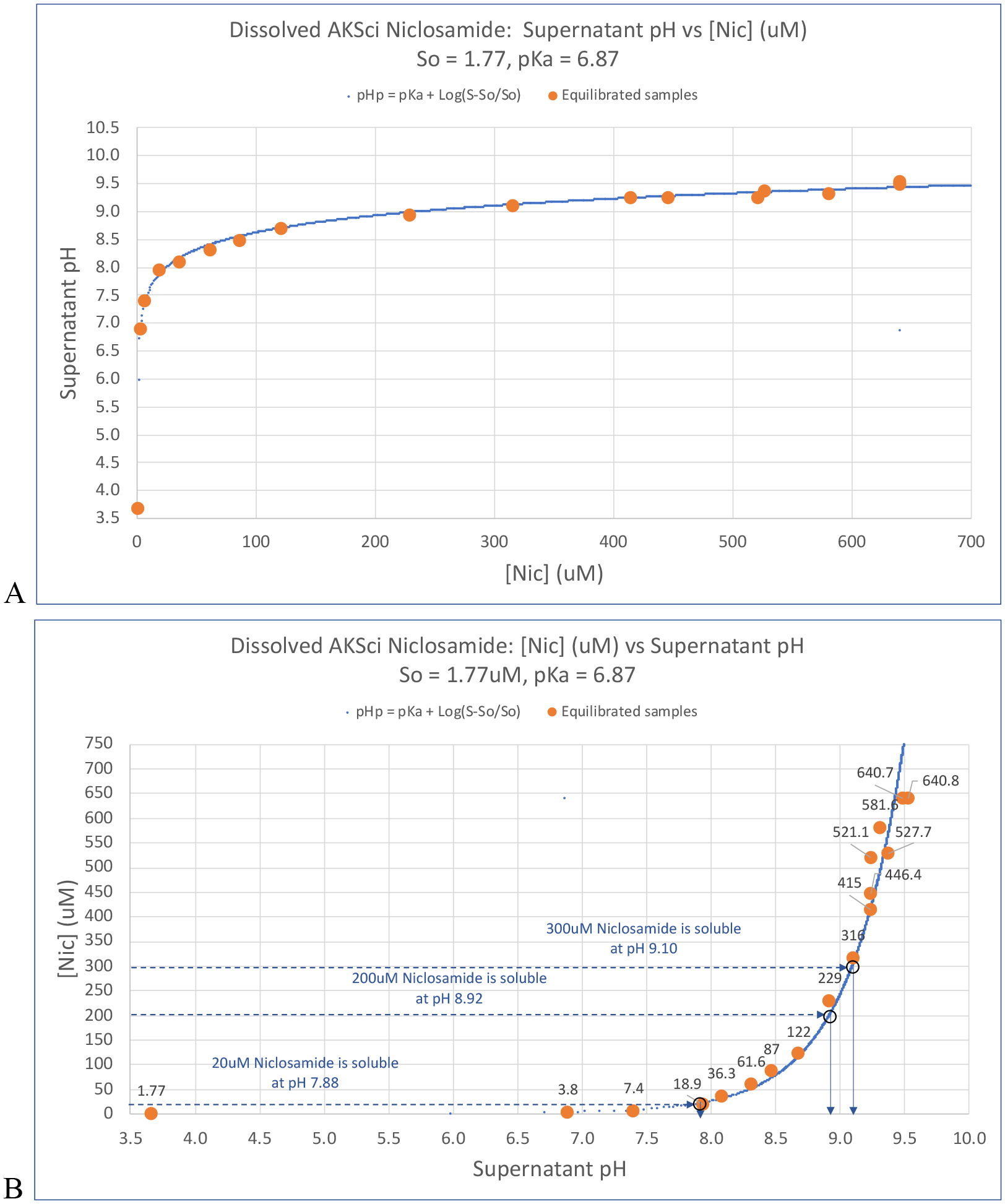
Equilibrated supernatant concentrations for dissolution of powdered niclosamide (from AK Sci) added in excess to each pH buffer. ***A)*** Supernatant pH versus Supernatant Niclosamide concentration [Nic] (uM) measured by nanodrop UV/Vis and compared to the pHp curve Eqn 4. Plotting this data against the pHp predictions for a measured S_o_ of 1.77uM, gives a fitted pKa for niclosamide of 6.87. ***B).*** Same data as in ***A*** but axes changed to a more easily evaluated form. Plotted is the Supernatant Nic concentration [Nic] (uM) versus supernatant pH. Again, also included is the pHp theory for pKa of 6.87 and limiting Nic_OH_ solubility of 1.77uM. As indicated by the dotted lines, a 20uM prophylactic solution can be made at pH 7.88; a 200uM early treatment throat spray can be made at pH 8.92, and the concentration can be raised to 300uM at pH 9.10. Limiting solubility of the deprotonated Nic_-ve_ is 640.7uM.

With increasing measured pH from 3.66 to 9.53 there is a concomitant increase in the supernatant concentration of niclosamide.

This data is well fitted in form and position by the pHp curve according to:

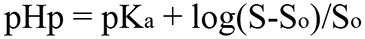

where, S_o_ = the molar solubility of the undissociated acid (Nic*_OH_*), and S = the molar concentration of the salt form in solution (Nic_-ve_). (See Eqn 4. ***Supplemental Information, S2.3***) using the measured S_o_ solubility of niclosamide at pH 3.66 of 1.77uM and a value for the pKa of 6.87.

In order to evaluate the data and compare to theory in what is, perhaps, a more easily evaluated form, the same data as in ***Figure 3A*** is replotted in ***Figure 3B*** with the axes switched. As can be seen, the supernatant niclosamide concentration ([Nic] uM) increased slowly over the lower pH range from 3.66 to just above 8, but then showed the expected more rapid rise in concentration from 8.5 to 9.5, where some of the highest values measured (and re-checked) for equilibrated samples were 641uM. This clearly demonstrates the potential for creating simple more concentrated solutions of niclosamide for the nasal and throat sprays.

The added horizontal lines in ***Figure 3B*** show that, for prophylactic use, a 20uM prophylactic solution can be made at pH 7.88, which is within the normal pH of the nasaopharynx (43). For the early treatment throat spray, as preclinical animal studies and then human studies proceed to dose escalation, a 200uM solution is readily achieved at pH 8.92, and the solution concentration of niclosamide can actually be raised to 300uM at only pH 9.10. In the oral cavity a higher pH is expected to be tolerated and this is where a higher niclosamide concentration is perhaps required for already infected epithelial cells and replicated and secreted lipid-coated virus particles embedded in the mucin secretions.

Comparing the pHp curve to the Henderson Hasselbalch curves (as in ***Supplemental Information, Figure S3***) shows that the data are actually consistent with the prediction of the amount in solution. For example, at the pKa where the amount of the acid in solution is measured to be 1.77μM, and so the total solubility at pH 6.87 should be 2 x 1.77uM = 3.54uM. The measured value at 6.89 is actually 3.8uM. Importantly for the nasal and throat spray application is that the amount of niclosamide in solution increases with increasing pH and was measured to be ∼641uM at pH 9.5 where, as shown in ***Supplemental Information Figure S3***, the dominant species is almost 100% Nic_-ve_. Therefore, these high pH values are expected to represent the solubility of that charged salt, i.e., 362 times more niclosamide is in solution at pH 9.5 than at pH 3.66, and this is all because of its pKa of 6.87 and a value for S_o_ of 1.77uM.

### 4.2 Rates of Dissolution for AK Sci Niclosamide powder

Having measured the equilibrium values of the amounts of niclosamide that can dissolve over a range of pH, the next series of experiments were to more carefully quantify the rates of dissolution of the AK Sci powder that was ground and suspended in buffer solution.

#### 4.2.1 Initial morphology of the AK Sci niclosamide powder

First, it is instructive to examine the initial morphology of the as-supplied AK Sci niclosamide powder. In ***Figure 4*** are photographic microscope images of AK Sci powder after grinding with a mortar and pestle, resuspending in water to disperse, and bath-sonicating to help break up aggregates for better visualization of individual particles.

**Figure 4.**
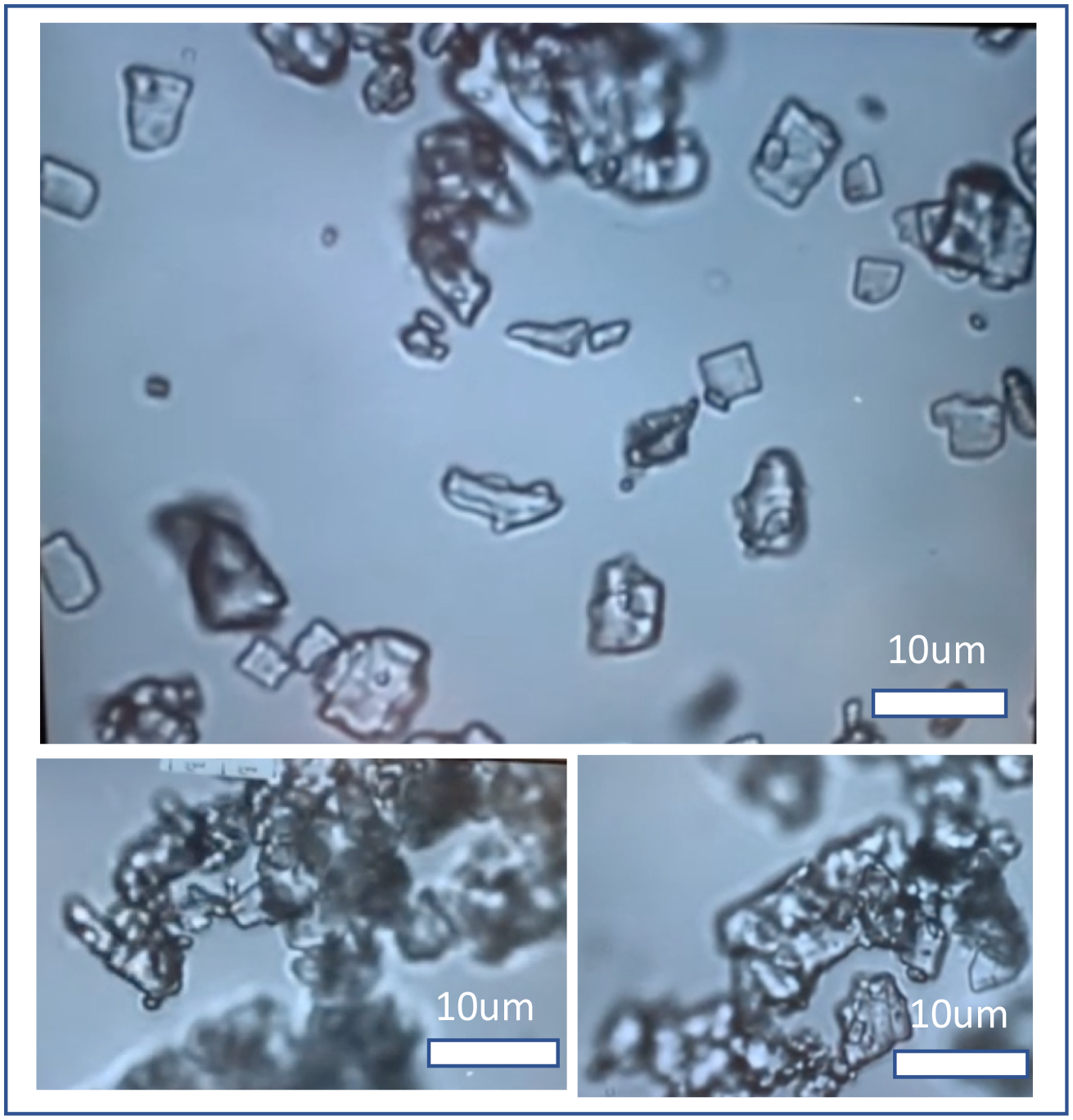
Optical microscope photographic microscopic images of AK Sci powder after grinding with a mortar and pestle, resuspending in water to disperse, and bath sonicating to help break up aggregates. Shown are typical particles that can be separate or in aggregated clumps of block-like material. (Bright field, 40x objective lens, with Köhler illumination).

Characteristic sizes, as length and width of individual particles, range from 1-10um with an average size (from this micrograph) of 4.3um +/-2.2um. Interestingly, any rod-like and spiky crystals that are characteristic of the low solubility monohydrate, (2) are noticeably and importantly absent. This is consistent with the AK Sci product being a solid form with a higher solubility than the stable monohydrate. While efforts were made to ask for details from AK Sci, suppliers were reluctant to divulge any information about product manufacture and post synthesis processing such as precipitation or recrystallization conditions, solvents, and potential solvates. Thus, the nature of the powdered material obtained from AK Sci remains, as yet, unknown.

Upon dissolution of the AK Sci material, i.e., when a portion of the added (1mM) niclosamide powder has dissolved and it was maintained under stirred conditions for 24 hrs and longer, subsequent microscopic observation showed that its morphology did not change radically from the original powder. Thus, the AK Sci niclosamide did not readily convert to the less soluble (spiky morphology) hydrates when incubated for days in the presence of its original powder. This observation is important to any drug formulations that utilize microcrystalline or even nanoparticles of niclosamide that are then in equilibrium with a suspending aqueous phase. That is, niclosamide from other suppliers, (e.g., Sigma) and when precipitated or made into “cosolvates” of water, ethanol and acetone, as presented and discussed below, each of these samples showed much lower solubilities (e.g. < 200uM at pH 9.3) characteristic of the most stable monohydrates and typical solvates investigated by van Tonder et al, (2, 60–62).

#### 4.2.2 Dissolution of AKSci (and Sigma) Niclosamide at the different pHs

While a more quantitative analysis would measure the surface area per gram of drug powder, all samples were ground and so somewhat standardized to a fine powder with the pestle and mortar (as shown in ***Figure 4)***. Therefore, for the same material (AK Sci niclosamide) the only assumed variables are total mass and mixing. Samples were weighed to within 10% of 3.27mg, which, in 10 mLs gives a total equivalent ∼1mM niclosamide, and so is in excess of all expected saturated supernatant concentrations. Since the mass added and mixing were similar for each pH sample tested, ***Figure 5*** shows that the Supernatant Niclosamide Concentration [Nic] (uM) versus time (mins) for the lower five dissolution curves gave a fairly quantitative measure of niclosamide dissolution at each pH of 8.62, 8.72, 9.06, 9.36, and 9.44.

**Figure 5.**
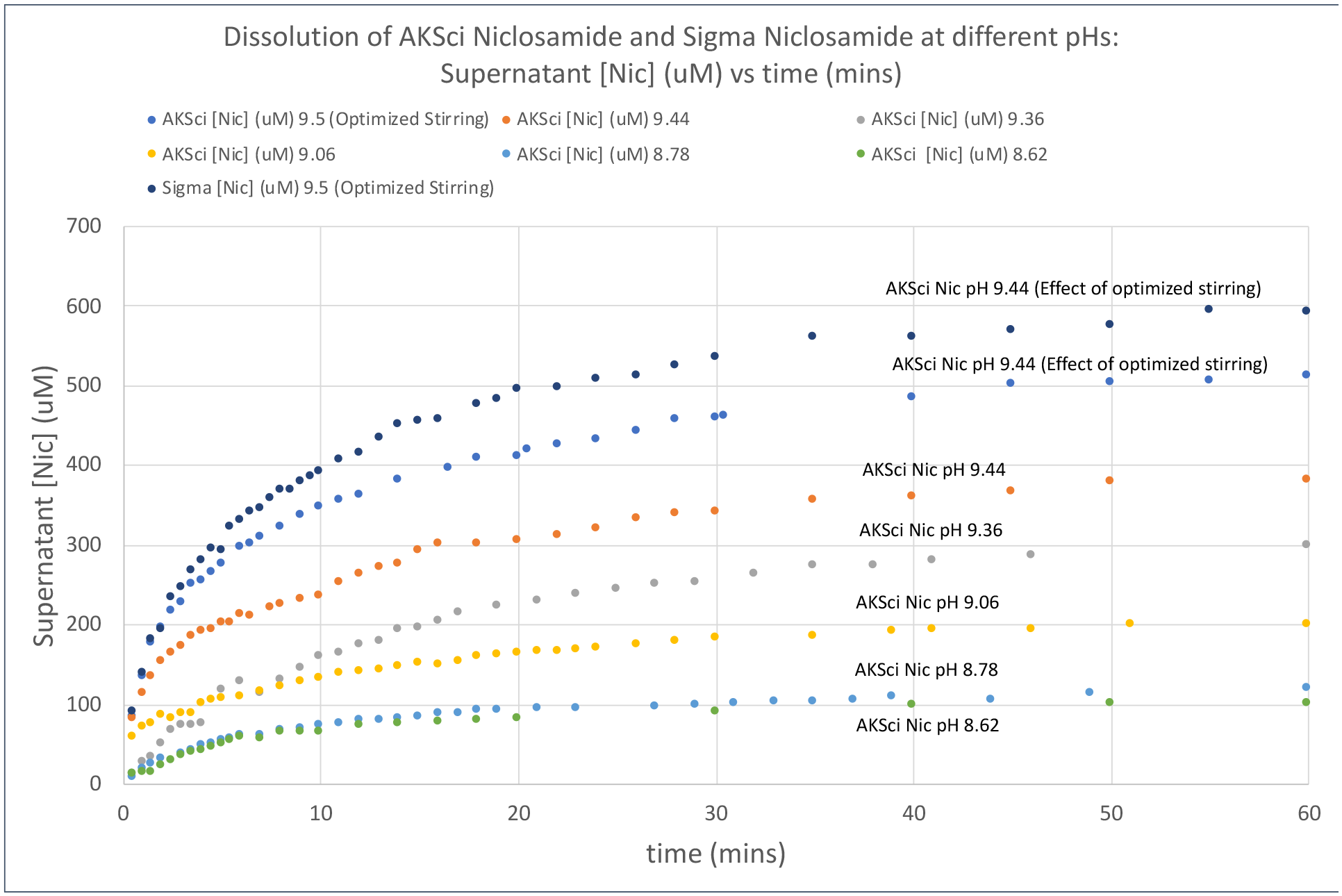
Dissolution curves for the different pHs over the first 1 hr after addition of dry powdered niclosamide. The lower five curves are dissolution data for the pH range 8.62 – 9.44 for a nominaltotal 1mM of the AK Sci material. The top two curves show the (empirical) effect of more optimized stirring, that increases the dissolution rate by ∼33%, for AK Sci and Sigma niclosamide.

As shown earlier in ***Figure 3B***, below pH 8.5, the equilibrium saturation concentrations are still quite low, and so we focused here on the higher pH range 8.62 – 9.5 for a nominal 1mM of the AK Sci material. The fitted logarithmic rate for each of these lower dissolution curves was found to increase with increasing pH as quantified by the pre-ln factor for the logarithmic fits for micromolar versus time in mins. These were, ***22.1*** (at pH 8.62); ***24.3*** (at pH 8.78); ***34.5*** (at pH 9.06); ***71.1*** (at pH 9.36) and ***69.0*** (at pH 9.44).

During these first experiments for pH 8.62 – 9.5, the magnetic stirbar was relatively small compared to the diameter of the scintillation vial, (about half) and the rotation speed was medium. In order to, at least empirically, explore the effect of stirring speed, a slightly larger magnetic stir bar was chosen such that the size of the magnetic stirbar was standardized to about 75% of the diameter of the vial keeping the stirring rate at medium. Under these conditions, the dissolution rate of niclosamide in pH 9.44 buffer was increased by about a factor of 37% i.e., to ***94.6ln(x)***. Also shown, the Sigma niclosamide dissolved slightly faster at ***113.8 ln(x)***. For all subsequent dissolution studies, this is the optimized stirring “standardized” set up.

#### 4.2.3 Initial dissolution rates

Dissolution was characterized above by the logarithmic fits, but it is more appropriate from a concentration per unit time perspective, to consider the initial dissolution rates over the first 3 min as intrinsic measures, i.e., intrinsic concentration units as micromoles of niclosamide dissolved per micrograms of niclosamide in the vial per second (uM/mg.s). ***Figure 6*** shows the initial (linear) dissolution rates derived from the data in ***Figure 5***, measured over the first 3 minutes, and plotted as a bar graph using the weight-based units.

**Figure 6.**
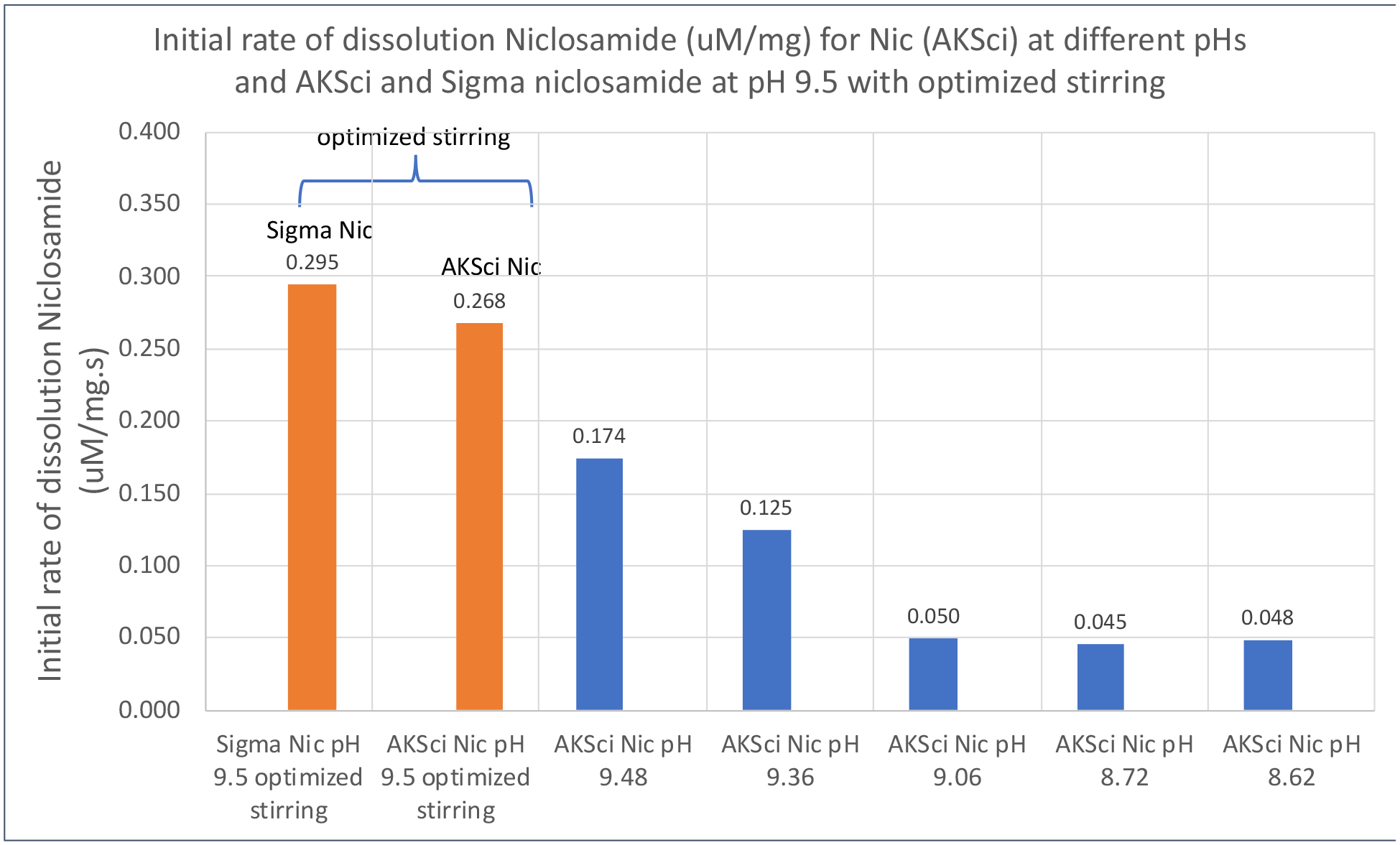
Initial rates for dissolution (uM/mg.s, i.e., uM of niclosamide dissolved per mg of niclosamide added to the vial per second) versus pH of the supernatant. Shown are the intrinsic concentration of niclosamide (uM/mg) over the first 3 mins of the plots in Figure 5 for AK Sci niclosamide into five different pH solutions of pH 8.62, 8.72, 9.06, 9.36 and 9.5. Bars in orange represent the effect of a more optimized stirring for AK Sci and Sigma niclosamide at pH 9.5

Consistent with the overall dissolution profiles, initial dissolution rates increased with increasing pH of the supernatant. Also shown is the effect of the more optimized stirring for AK Sci at pH 9.44 giving an increased rate of dissolution as expected from dissolution models. This more optimized stirring increased the initial rate of particle dissolution by a factor of about 50% but did not change the final equilibrium saturation concentration. Also shown, the dissolution rate for the Sigma sample is similar the AKSci sample.

Dissolution models (like Epstein-Plesset (65) that we have used previously to evaluate dissolution of gas (66, 67) and liquid microparticles under diffusion controlled conditions (68–70)), assume that the concentration at the particle surface is the saturation concentration C_s_, and that the rate of dissolution (dm/dt) should be proportional to this saturation concentration. Also included is the thickness of the stationary layer that, itself is a function of stirring. Thus, as shown in ***Figure 7***, when plotted as the normalized concentrations (uM/mg) the initial rates of dissolution (uM/mg.s) are close to being proportional to the equilibrium solubilities, especially for the higher pHs and hence higher amounts in solution from ***Figure 3B*** and the AKSci and Sigma niclosamide (where the stirring was standardized).

**Figure 7.**
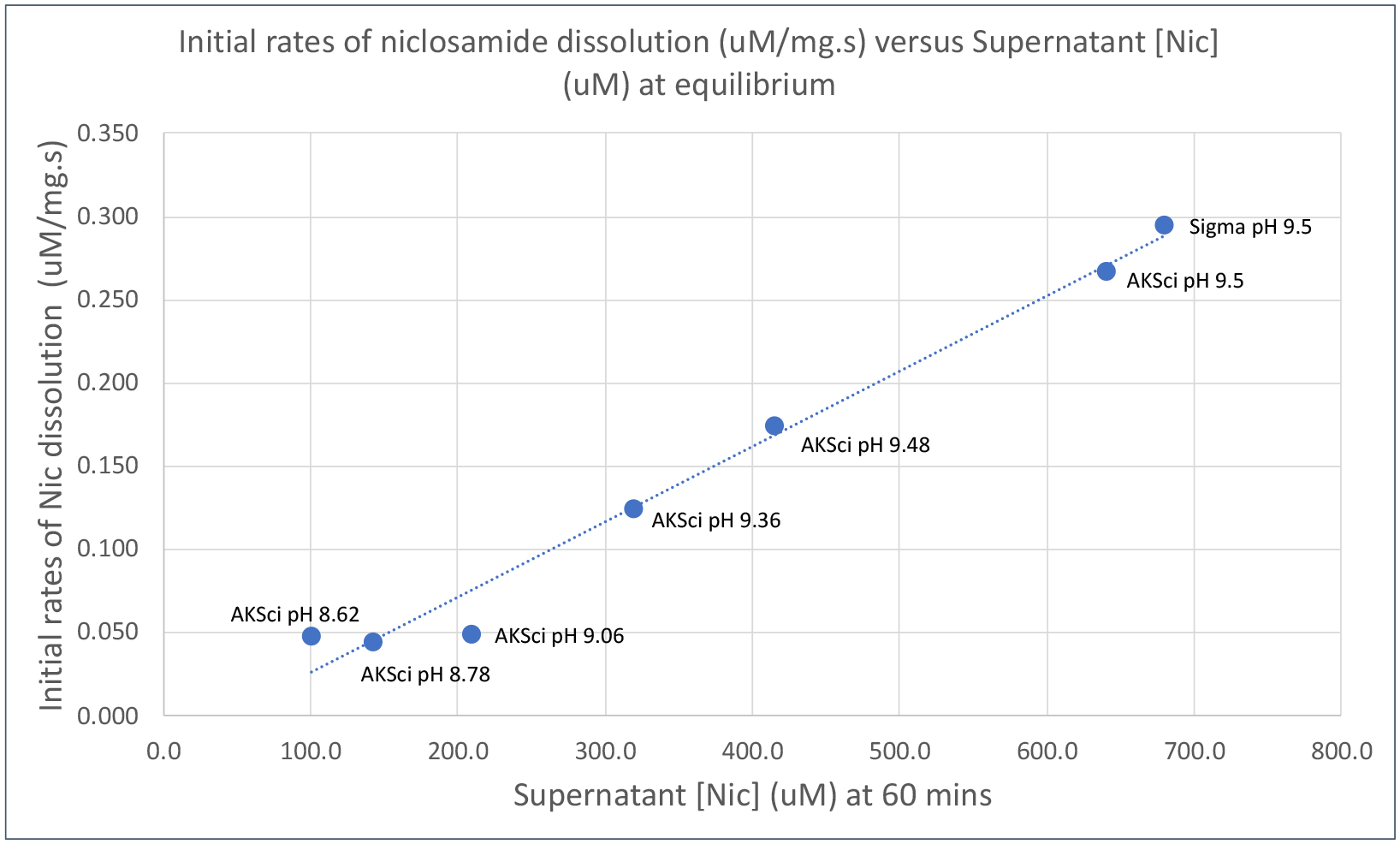
Initial rates of niclosamide dissolution Supernatant [Nic] (μM/mg.s) versus Supernatant niclosamide concentration [Nic] (μM) at equilibrium (from the data in Figure 3B) and the Sigma value.

### 4.3 Supersaturated Solution precipitation and their “natural” morphologies

#### 4.3.1 Supersaturated precipitation compared to the dissolved AK Sci Niclosamide versus pH

As used to accurately make the control solutions, the second way to achieve a solution of niclosamide is to first dissolve the niclosamide in a water-miscible solvent (like ethanol, or acetone, DMSO, or DMA) and exchange the solvent for the aqueous anti-solvent at a final concentration where the niclosamide is still soluble (see above, 4.1.2 Solvent exchange technique). This technique can also be used to create supersaturated solutions from which the niclosamide can (eventually) precipitate and form whatever the stable solid morphology is at that pH. Thus, in these preliminary experiments, levels of supersaturation were explored and compared to the parent AK Sci material, including the final equilibrium solubilities of the precipitated niclosamide as a function of the supernatant pH. Niclosamide was formed into supersaturated solution over the range of pH’s at 2-5 times excess niclosamide (with respect to the equilibrium amount of AK Sci niclosamide in solution as in ***Figure 3B*** and ***Table 1***) and allowed to precipitate. For comparison, shown in blue symbols, is a series of AK Sci niclosamide dissolved from ground-powder, as given earlier in ***Figure 3B***. The orange symbols are niclosamide from the same source(AK Sci) that was first dissolved into ethanol at 30mM and then injected into each buffer at 2-5 times the excess concentration of that measured solubility, allowed to precipitate, and fully equilibrate for 8 days and filtered through a 0.22µm filter prior to taking the clear supernatant for measurement. Thus, as shown in ***Figure 8***, the precipitated niclosamide has much lower supernatant niclosamide concentration than the parent compound, that is simply dissolved in each pH buffer.

**Figure 8.**
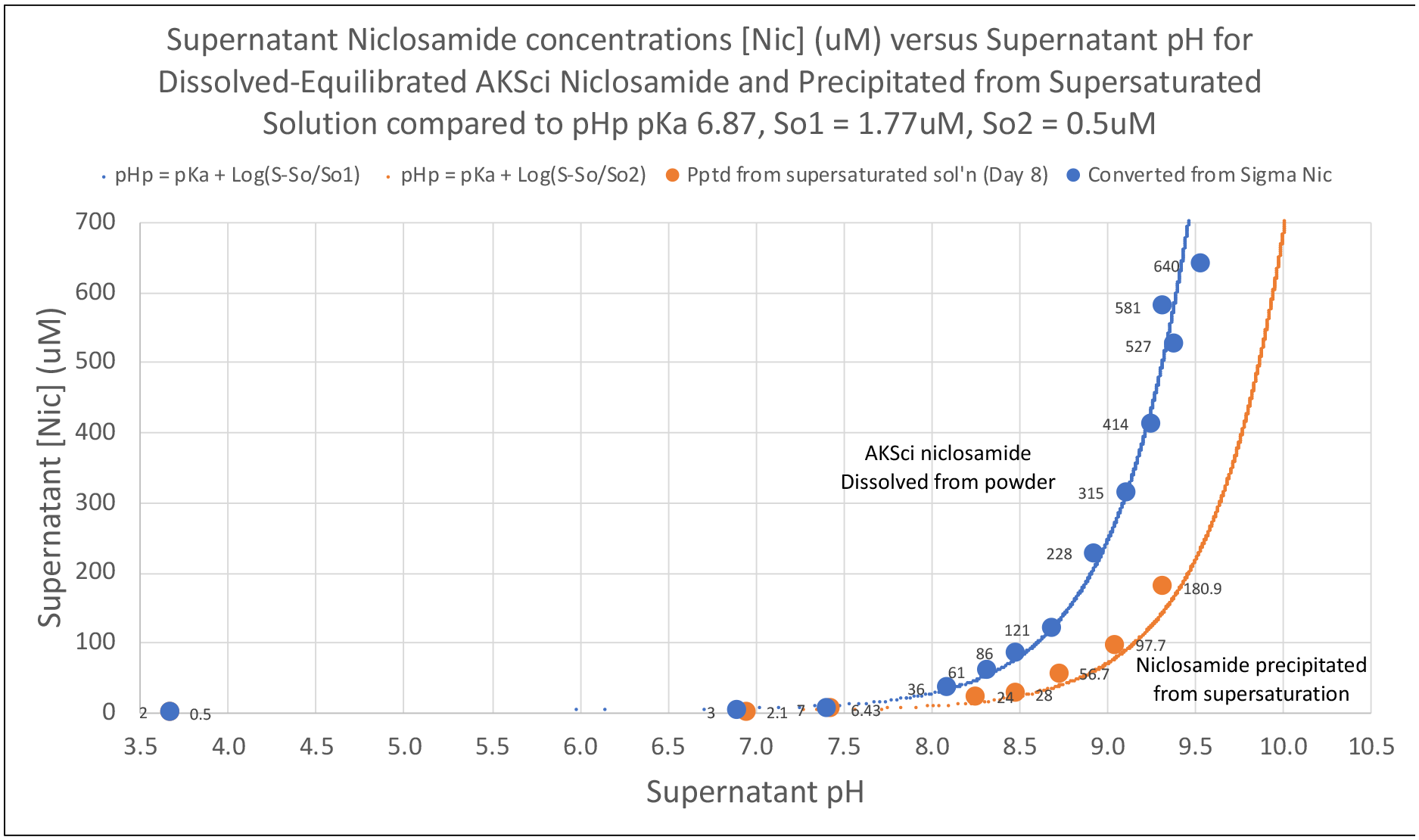
Supernatant Niclosamide concentrations [Nic] (mM) versus supernatant pH for dissolved-equilibrated AK Sci Niclosamide in solution (blue symbols) and precipitated from supersaturated solution (orange symbols) compared to pHp with pKa = 6.87, So1 (AK Sci dissolved) = 1.77uM (blue line); S_o_2 (precipitated from supersaturation) = 0.5uM (orange line).

The blue and orange lines are the pHp curves for each system where the pKa is the same one (6.87) derived from fitting the AK Sci dissolution data from ***Figure 3A*** and ***B***. The measured value of the solubility of the fully protonated acid for the parent AK Sci niclosamide, labelled here, as S_o_1 of 1.77uM ± 1.0uM (blue curve) is used along with the measured equilibrium supernatant niclosamide concentration for the precipitated niclosamide (orange curve), S_o_2, of 0.5uM ± 1.0uM measured at pH 3.66; this represents the limiting S_o_ solubility of the protonated acid made after precipitation in equilibrium with its precipitate at that pH. Since the pKa is the equilibrium balance between the acid and salt forms in solution, it is satisfying that the same pKa of 6.87 can be used to fit both sets of data when the measured value for the solubility of the precipitated Nic_OH_ is also used. There was some difficulty in measuring these low pH solubilities, but there was a discernable difference between instrument noise for the blank and the sample, hence a plus minus of 1uM. If we use a value for S_o_2 of 0.6uM for precipitated niclosamide at pH 3.66 this actually fits the data even better and is within the standard deviation of this difficult-to-obtain value.

What this data shows is that niclosamide, when precipitated from supersaturated solution, equilibrates to a much lower final supernatant niclosamide concentration, which is presumably reflective a more stable polymorph at each pH. Prior to precipitation, it was possible to achieve supernatant concentrations (kinetic solubility) as high as 3mM niclosamide at the highest pH of 9.5, -a supersaturation of ∼ 3mM/200uM = 15 times. The data shows that, while the AK Sci niclosamide powder was relatively stable in solution at the high concentrations achieved, precipitation from supersaturated solution presumably forms the water-hydrate at each pH, that still does have a pH dependence for its now thermodynamic solubility.

One question here might be, “Could the 1% ethanol that was introduced into the buffer during making by solvent exchange be the source of the difference, not the hypothesized difference in crystalline form?” First, morphologically speaking the images of the AK Sci powdered niclosamide particles that are in equilibrium with the supernatant solutions are quite different from the precipitated particles. At pH 9.5 for the dissolved sample, they still look like the block-like particles shown in ***Figure 4***, whereas the precipitated particles that were made by solvent exchange at pH 9.3 show the spiky appearance as in ***Figure 9***. Also, in the case of niclosamide, its solubility in ethanol is ∼35mM (measured here, data not previously shown) and its solubility in pH 9.5 buffer is 641uM. As mentioned earlier (3.2.1) the Yalkowski model for cosolvent solubility (71–73), shows that if the compound is more soluble in ethanol than in water, then in a water-ethanol mixture the solubility of the compound increases with increasing ethanol, not decreases. However, the data in ***Figure 8*** shows a decrease in the amount of niclosamide in solution in the 1% ethanol, (that is still 273mM). This is also in line with the niclosamide monohydrate solubilities reported by van Tonder (2) and seen for the “cosolvate” recrystallized from ethanol. Future studies that characterize the pH dependent and cosolvate dependent crystal structures will confirm or modify these hypotheses.

**Figure 9.**
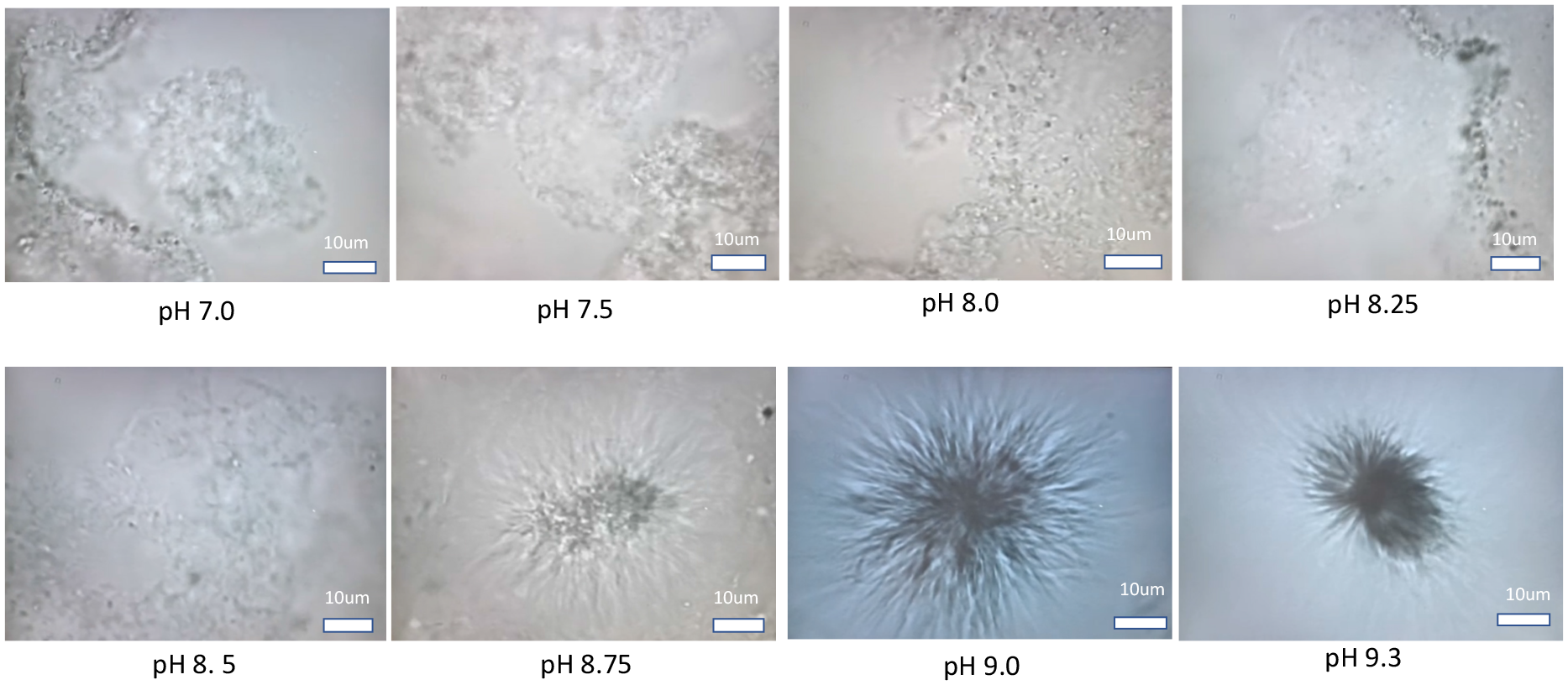
When precipitated from supersaturated solutions AK Sci niclosamide forms a series of particle morphologies of reduced solubility such that the amount in a solution that changes with pH. Shown are optical microscope images (bright field, 40x objective lens, with Köhler illumination) taken from the precipitated samples at each indicated pH. Scale bar is 10um.

#### 4.3.2 Supersaturated solutions form a series of niclosamide morphologies of reduced solubility

Corresponding to the supernatant solubilities given in ***Figure 8***, samples of the precipitated material were taken and viewed under the optical microscope. Shown in ***Figure 9*** are the photographic images of precipitated particles at each supernatant pH as full screen images using a 40x bright field objective.

At pH 7, where the low solubility protonated acid makes up almost 50% of the niclosamide in solution (see Henderson Hasslebalch curve in ***Supplemental Information***, ***Figure S3***), niclosamide precipitates as flat particulate sheets formed in the stirred aqueous buffer that make a mass of gel-like particles in suspension. Macroscopically, these are the characteristic white particles seen swirling in the vial (in ***Supplemental Information***, ***Figure S1***). This flat sheet, particulate, gel-like morphology persists at pH 7.5, 8.0, 8.25 and 8.5. Being formed from a 1:99 dilution of ethanolic niclosamide that is exchanged for the excess pH buffer under rapid stirring, some sheets are observed to fold, as in the image for pH 8.25. Then, at pH 8.75, the morphology makes a transition to the more usual monohydrate spiky polymorph that is characteristic of the precipitated particles at pH 9.0 and 9.3, where the deprotonated niclosamide salt is the dominant species. What these microscopic images demonstrate then, seemingly for the first time, is that these most stable niclosamide hydrates (2) not only have a pH dependence to the amount of niclosamide in solubility but also have a pH dependent morphology.

#### *4.3.3.* Niclosamide gel-like particles also display a strong hydrophobicity and coat gas bubbles

As shown in ***Figure 10***, in some images of the samples of the precipitates dark structures were often observed. This is a new and interesting, but not unexpected, observation that the niclosamide gel-like particles also displayed a strong hydrophobicity. What these represent are gas microbubbles that follow the crumpled contours of the niclosamide precipitated sheet formed at pHs 7.0, 7.5, 8.0 and 8.5 as the precipitate adheres to the bubble surface. Because the particles are formed by precipitation in a rapidly stirred buffer environment, it seems that gas microbubbles can get trapped. These characteristically optically black masses of air show a complete lack of surface tension (or tension in the surface). That is, rather than being round and exhibiting their usual air-water surface tension and concomitant Laplace pressure, they are deformed to the shape of the niclosamide material. Clearly, as one would expect from niclosamide’s low solubility and moderate logP of 4, at these pHs, the niclosamide particles are quite hydrophobic. This was also borne out by the way the particles of powder, when added to a vial for the dissolution test, would rapidly float at the air solution interface unless well shaken to wet the powdered particles and immerse them in buffer suspension.

**Figure 10.**
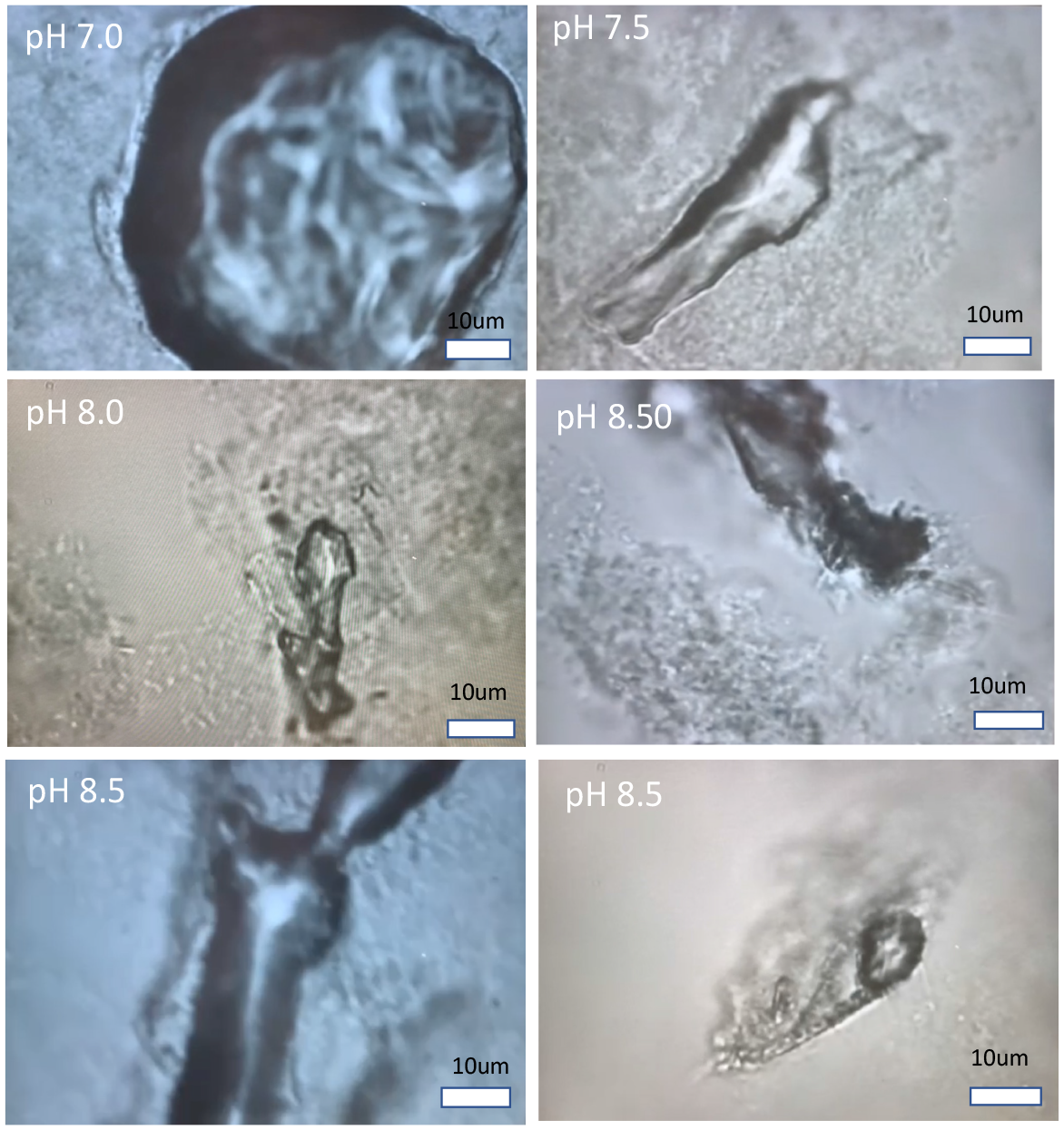
The niclosamide gel-like particles also display a strong hydrophobicity. Gas bubbles, that were adsorbed or trapped in the microparticles due to stirring during initial mixing and subsequent stirred equilibration, appeared to follow the crumpled contours of the niclosamide precipitated sheet formed at pHs 7.0, 7.5, 8.0 and 8.5. The precipitate adhered to the bubble surface showing that the interfacial tension and Laplace pressure were zero. (Bright field, 40x objective lens, with Köhler illumination).

This kind of zero tension and zero Laplace pressure has actually been observed and well-characterized before for lipid coated gas microbubbles (as solid lipid monolayers) in our micropipette experiments that measured interfacial tensions as well as gas bubble dissolution in undersaturated solutions (66, 74, 75). Here though, the hydrophobic precipitated niclosamide sheets form a kind of “Pickering gas-emulsion” that is stabilized by the sheet-like particles adsorbed onto the interface between the aqueous and gas phases.

### 4.4 Dissolution of Niclosamide as a water-precipitate, and as recrystallized material from Acetone and Ethanol

#### 4.4.1 Dissolution profiles

Given the high solubilities observed by the AK Sci material, while other sources and precipitated material readily converted to the less soluble and more stable hydrates, preliminary studies were conducted to create different precipitates and cosolvates in order to evaluate their dissolution and crystal morphologies. These studies were conducted to initially compare the samples with the AK Sci and Sigma materials and try to ascertain if they were a water, ethanol or acetone solvate; they did not appear to be so.

Samples of niclosamide were made from the AK Sci original material as a water-precipitate and recrystallized from Acetone and Ethanol, as described in methods broadly following those by van Tonder et al for their well-characterized cosolvates (2, 60, 62). The same mass of ground material, 3.5mgs, was added to each 10mls of pH 9.3 buffer in a 20mL scintillation vial and stirred with magnetic stir bar. 2uL samples were taken at time intervals and their absorbance measured on a UV/VIS nanodrop spectrophotometer at 333nm. The concentrations of niclosamide (uM) in the supernatant versus time (mins) are plotted in ***Figure 11*** for the water precipitate, and the recrystallized niclosamide from acetone and ethanol. Dissolution profiles were therefore determined for each of the dried samples (water precipitated, and acetone and ethanol recrystallized) as well as the equilibrium supernatant concentrations several days after dissolution, all determined at pH 9.3. As can be seen, the dissolution rates go as Nic_H20ppt_ > Nic_Acetone recrys_ > Nic_EtOH recrys_, although the initial rates measured as the intrinsic parameter uM/mg of material were: ^Nic^_H20ppt_ ^(0.^104 uM/mg) > Nic_EtOH-recrys_, (0.0439uM/mg) ∼ Nic_Acetone-recrys_ (0.0369uM/mg).

**Figure 11.**
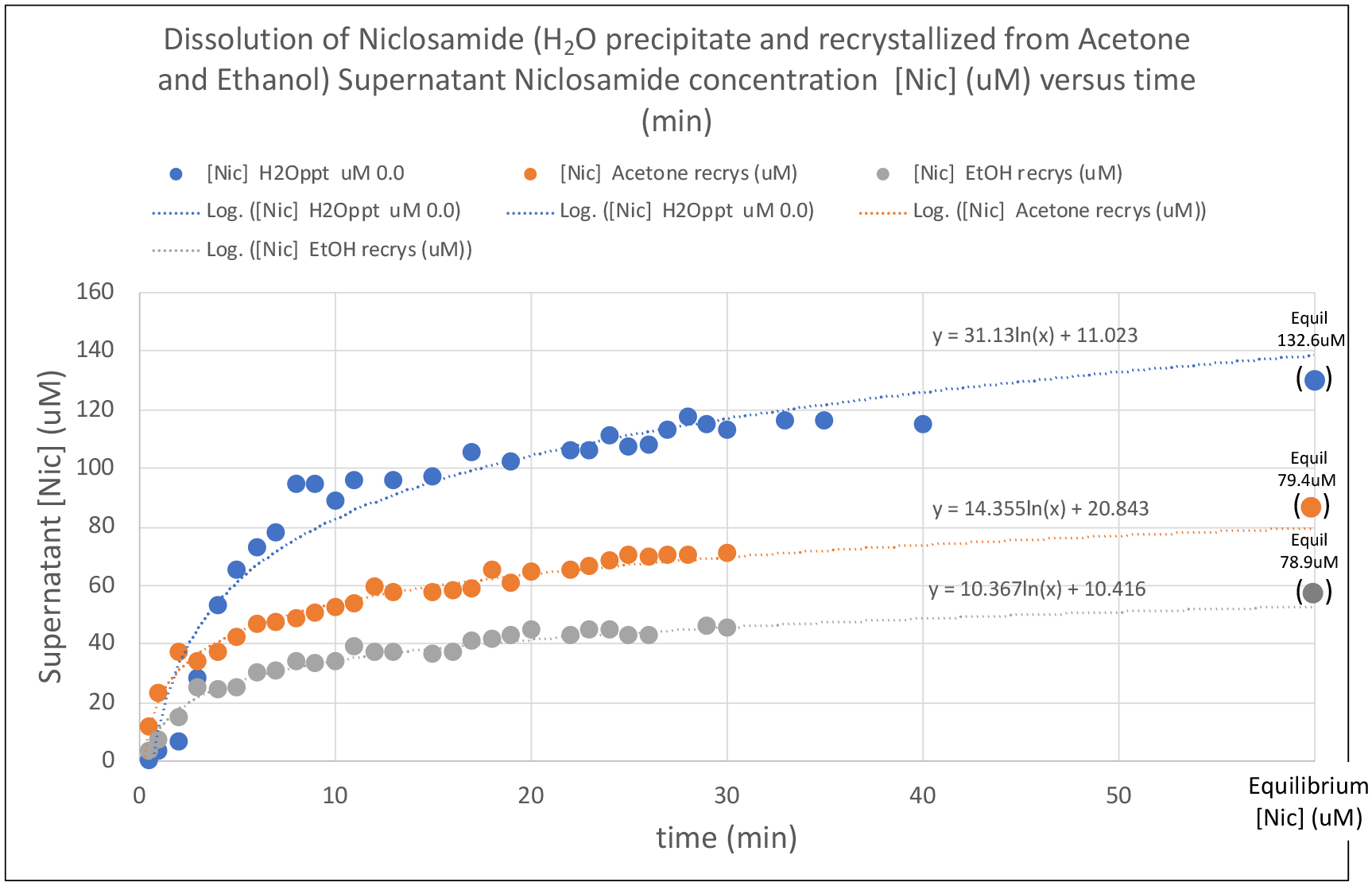
Dissolution of Niclosamide (H_2_O precipitate and recrystallized from Acetone and Ethanol) Supernatant Niclosamide concentration [Nic] (uM) versus time (min). Also shown are the overall rates and final equilibrium supernatant concentrations of niclosamide several days after dissolution was started.

This data shows that the commercial niclosamide products from AK Sci and Sigma were likely not recrystallized from acetone (as we were told) because the as-received AK Sci and Sigma niclosamide samples attained a much higher concentration in solution (∼550uM) at pH 9.3 than the acetone recrystallized material (79.4uM). It also shows how the water “solvate” that was dried and then re dissolved at pH 9.3 has a similar (132.6uM) solubility to that precipitated *in situ* as in ***Figure 8*** of 180.9uM. Also, the acetone and ethanol (presumed) cosolvates have an even lower solubility than the water precipitated material.

#### 4.4.2 Crystal morphologies

Images of the morphologies of the water-precipitated and acetone- and ethanol-recrystallized samples are shown in ***Figure 12***.

**Figure 12.**
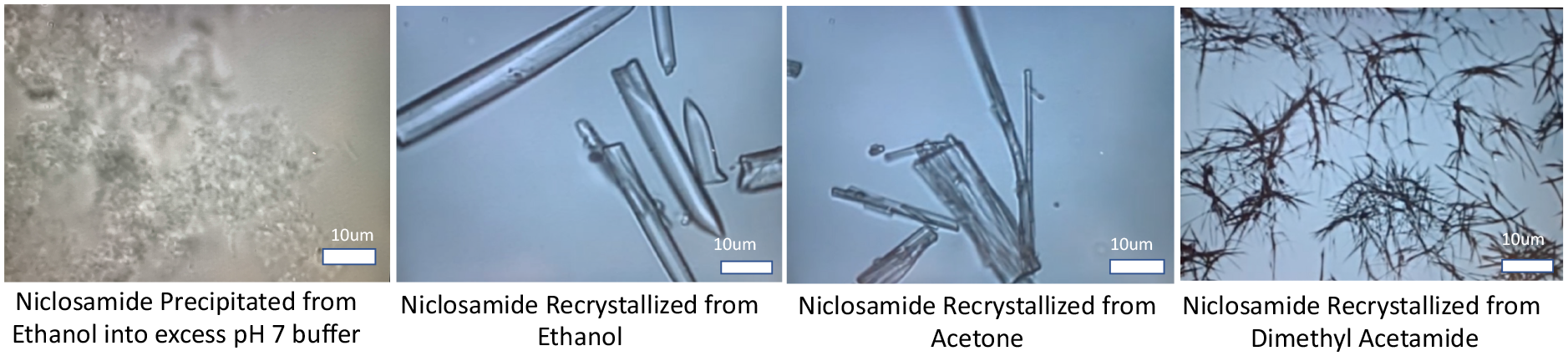
Optical microscope images of Niclosamide precipitated from supersaturation (at 1%ethanol), and Niclosamide recrystallized from ethanol, acetone and dimethyl acetamide. (Bright field, 40x objectivelens, with Köhler illumination).

As shown earlier (***Figure 9***), and included here for comparison, niclosamide precipitated from supersaturated solution (after solvent exchange from an ethanolic solution into excess water at 1% ethanol) has a very different morphology to the rod-like samples recrystallized from acetone and ethanol. These latter structures are more reminiscent of the methanol and other cosolvates made and characterized by van Tonder et al (2, 60–62). Additionally, although not measured in dissolution studies, is an image of a sample of niclosamide recrystallized from dimethyl acetamide that had a finer and more fibrous morphology.

These images taken of the various precipitated and solvent-recrystallized samples illustrate that the source of niclosamide, including its synthesis and post processing solvent recrystallization or precipitation recovery, can dramatically influence the final solubility of the drug compound. This, in turn, is expected to influence the pharmaceutical performance of the drug product, especially its solubility limit and dissolution, both of which affect its ultimate bioavailability. This is especially important in our (and other’s) applications as nasal and throat sprays. As mentioned above, the most effective way for niclosamide to permeate through the mucin layer that normally covers and protects the underlying nasal and buccal epithelial cells is as a soluble molecule. Also, any suspension of microparticle material is expected to undergo a conversion to the more stable and least soluble monohydrate form. However, if the solid is filtered out or removed by centrifugation, the remaining supernatant retains its high soluble form, unless heterogeneous nucleation might trigger the monohydrate.

### *4.5* Sigma Niclosamide dissolution and overnight stirring also produces lower solubility material

As shown above (***Figure 5***), the Sigma Niclosamide product readily dissolves and, over a period of 3 hrs, it achieved a niclosamide supernatant concentration of 680uM. While slightly more soluble (AK Sci was 640uM) this is very much in line with the AK Sci material at this pH. However, while the AK Sci material appeared to be generally stable in the presence of the excess powdered particles that were in equilibrium with their supernatant solution, as shown in ***Figure 13***, the Sigma material converted to a much lower solubility solid form over a period of the next several hours. 24hrs after initially starting the dissolution experiment, as shown in ***Figure 13***, the supernatant went from a clear solution (top image) to a cloudy suspension (lower image). When the supernatants were filtered through 0.22um filters and the supernatant niclosamide concentration was measured by UV/Vis spectroscopy, the supernatant niclosamide concentration had fallen to 198uM at ∼pH 9.6, --a loss of material concentration of 482uM.

**Figure 13.**
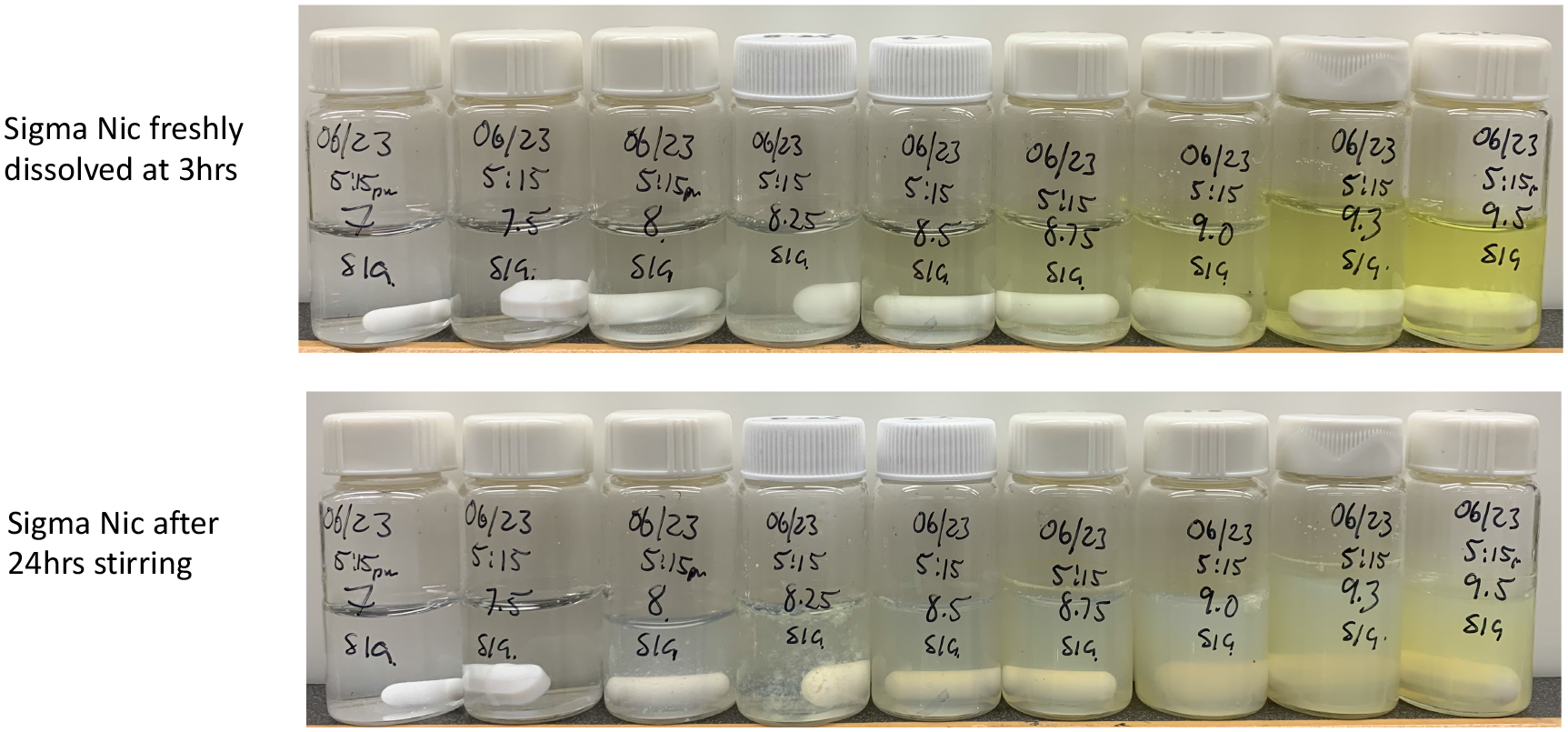
Comparison between (top) Sigma niclosamide freshly dissolved at 3hrs, and (bottom) after 24hrs stirring, showing the clear supernatant converted into a cloudy suspension. The supernatant niclosamide concentration was also decreased as shown in Figure 14.

Thus, as shown in ***Figure 14***, when dissolved and equilibrated over the range of pHs 7 to 9.5, the final supernatant concentrations were in a similar range to those measured for niclosamide precipitated from supersaturated solution (in ***Figure 8***). i.e., they both showed solubilities (SigmaNic at pH 9.3 = 103uM; precipitated material = 180.9uM) consistent with forming the more stable hydrated material. Again, comparing the experimental data to the pHp model using the same derived pKa of 6.78, for the fit a value of S_o_ is used to fit the data that is again consistently low, ∼0.35uM.

**Figure 14.**
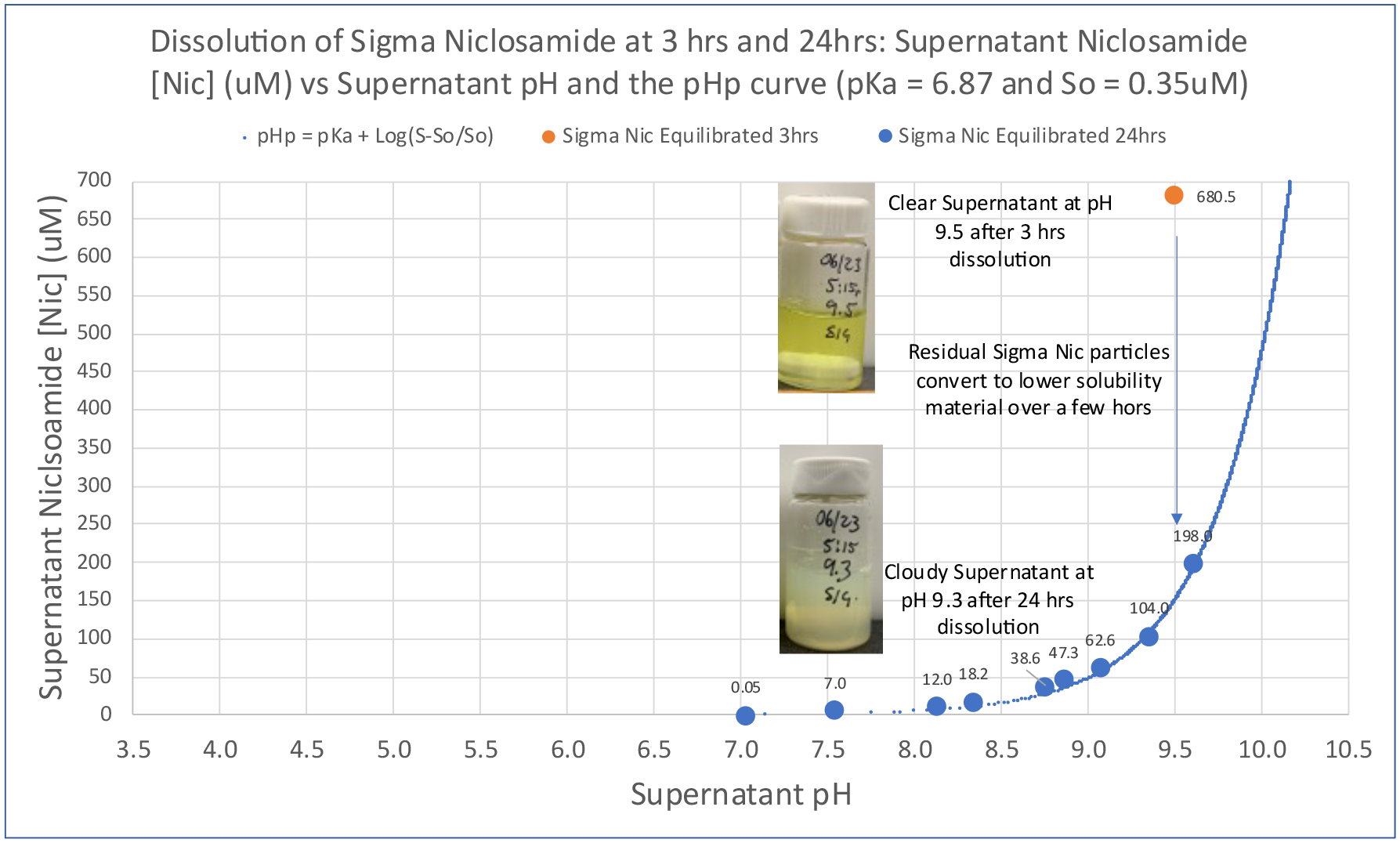
Dissolution of Sigma Niclosamide in pH 9.3 buffer at 3 hrs and 24hrs: Supernatant Niclosamide [Nic] (uM) vs Supernatant pH and the pHp curve (fitted with pK_a_ = 6.87 and S_o_ = 0.35uM). Also shown are photographic images of the same 10mLs of suspension in the 20mL scintillation vial for the clear supernatant after 3 hr dissolution and the much cloudier overnight stirred sample.

Finally, as shown in ***Figure 15***, this overnight equilibration and resulting decrease in supernatant concentration produced a change in particle morphology. The block-and-rod-like particles of the original Sigma niclosamide material were converted to long, more fibrous, and spiky bundled structures with a background of smaller fainter fibers that have a much lower (∼3.5x) solubility than the parent Sigma niclosamide powder. It is interesting to note that the difference between the Sigma niclosamide product and the AK Sci product (***Figure 3***) is that, while both have an appearance of block like crystals, the as-supplied Sigma material does have a rods and spike morphology, and so were likely processed after synthesis by recrystallization or precipitation in different media. Also, the powdered material is more prone to conversion to the more stable (apparent) hydrate after overnight equilibration in the pH 9.3 buffer.

**Figure 15.**
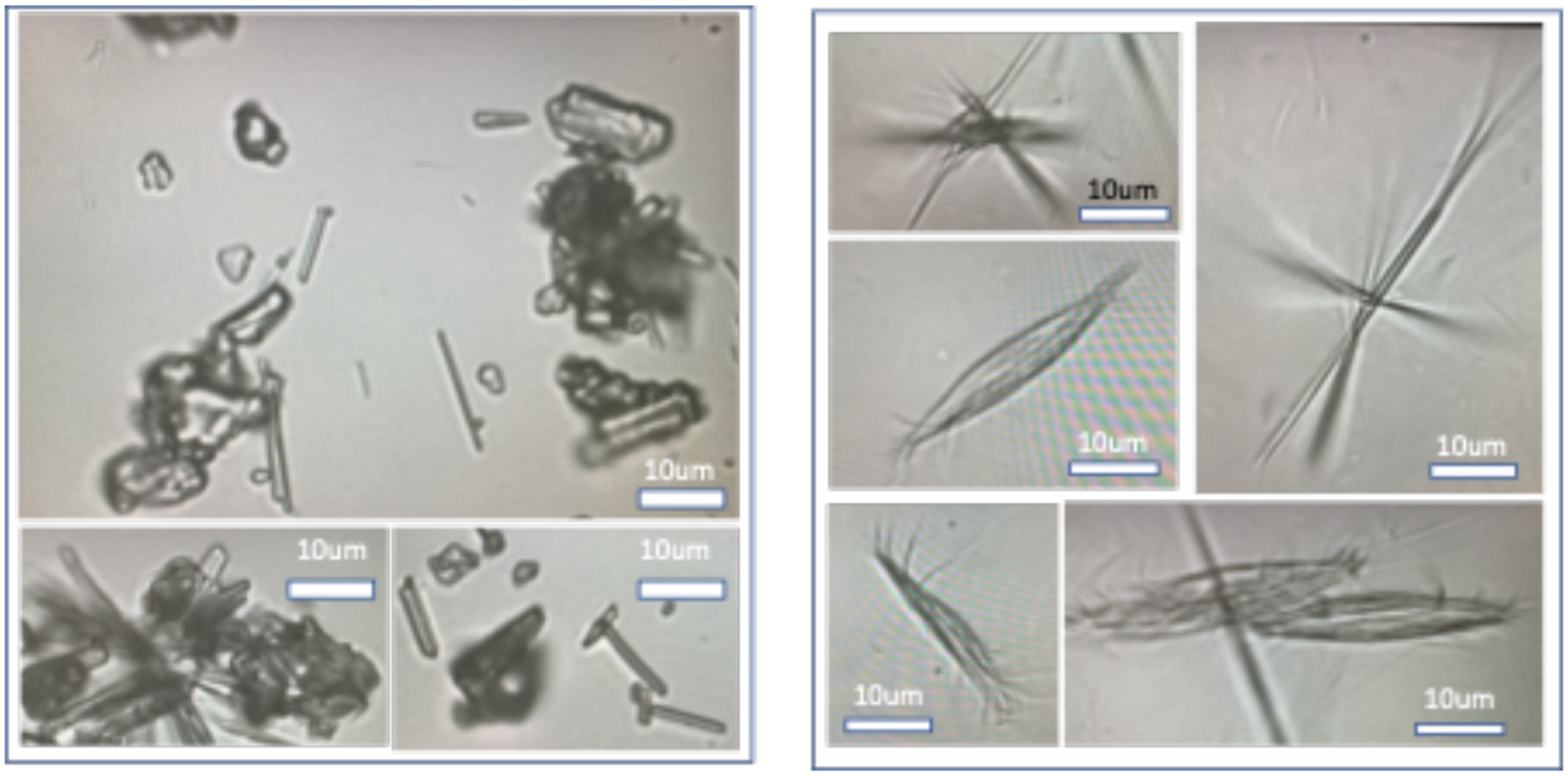
Optical microscope images of (left) the Sigma Niclosamide as received from the supplier and (right) after overnight equilibration at pH 9.3. Overnight equilibration of Sigma Niclosamide (right) produced long rod-like and spiky-bundled structures and a background of smaller fainter rods that have a much lower (∼3.5x) solubility than the parent Niclosamide powder.

All of the above niclosamide particles (AK Sci, Sigma as supplied, precipitated, converted and recrystallized) would now require crystallographic analyses for measuring polymorphic, hydrate, and amorphous states as could be done by x-ray crystallography and/or FT-IR/Raman, as well as calorimetric evaluation, as used here for two polymorphic forms of sulindac with very different solubilities (76).

## 5. Discussion

The main results of this study are that (1) Niclosamide has a pH-dependent solubility and dissolution rate, and (2) solubility and the amount of niclosamide in supernatant solution is determined by the nature of the solid form in contact with the supernatant solution and ultimately the most thermodynamically stable crystalline forms. Here, “solubility” refers to the actual solubility of the two niclosamide species, i.e., the protonated acid (Nic_OH_) and unprotonated salt (Nic_-ve_) which are measured to be 1.77uM (at pH 3.66) and 641uM at pH 9.5. Between these two pHs the important quantity is the amount of niclosamide actually in solution. This is therefore given by the 1.77uM of the acid (at its solubility limit) plus the amount of the unprotonated salt that increases with increasing pH.

Thus, the picture that emerges from this series of studies is that the amount of niclosamide in aqueous solution follows the predicted pH dependence from the Henderson Hasselbalch and pHp models. The reason these results are important is that niclosamide is currently being considered, formulated, and in some cases tested in clinical trials in a range of formulations, including a micronized niclosamide in spray dried lysozyme (37, 38, 77, 78). Given the poor solubility of niclosamide at neutral and lower pH as expected in the nasal pharynx, understanding, and more fully characterizing the pH dependence of niclosamide in aqueous buffers therefore allows a more optimized simple solution to be used for nasal and throat sprays.

Also, niclosamide is able to take on different morphological forms as already established to some extent by van Tonder et al (2, 60–62), but now showing this for niclosamide from different suppliers, and over a range of pHs. As is well known in pharmaceutics, although identical in chemical composition, polymorphs differ in bioavailability, solubility, dissolution rate, chemical and physical stability and may other properties (76). As encouraged by Llinàs et al (76), *“Despite significant investment in processes to find all the possible polymorphs of active pharmaceutical ingredients (APIs), new polymorphs can suddenly appear without warning. Polymorphs tend to convert spontaneously from less stable to more stable forms, and, therefore, it is best to discover and characterize the stable form as early as possible”*

Thus, any new formulations of niclosamide that are sought for testing in cells, animals, and especially in humans, and that are expected to deliver niclosamide, need to take into account the ultimate solubility of any more stable, in this case, monohydrate forms (2). If they occur, or are allowed to occur, they will determine the amount of bioavailable niclosamide that can permeate the mucin in a nasal spray or inhaled administration. In our niclosamide solution approach they are not present and so cannot exert any deleterious equilibrating effects, i.e., the high niclosamide solution concentration is preserved.

Discussed next are the main experimental results associated with the equilibrium and kinetic dissolution experiments for niclosamide into pH buffers from commercial sources and how this enables making the niclosamide-based nasal and throat sprays. This is followed by additional scientific findings that when precipitated by solvent exchange or when supernatant solutions are left in contact with the excess solid material, niclosamide readily reverts to more stable and lower solubility polymorphs. Filtering out the excess niclosamide preserves the achieved amount in solution.

### 5.1 Niclosamide solutions can readily be obtained by dissolving niclosamide powder into simple buffer

As predicted in ***Figure 4***, and measured in ***Figures 7A*** and ***7B***, a 20uM niclosamide solution can readily be obtained by dissolving niclosamide powder into aqueous solution at a pH of 7.88; a 200uM solution of niclosamide can be obtained in buffered solution at pH 8.92; and the concentration can be raised to 300uM at pH 9.10. The ultimate solubility of the negatively charged niclosamide salt (Nic_-ve_) at pH 9.5 was 641uM.

For the nasal epithelium, while the average pH in the anterior (front) of the nose is 6.40 and the average pH in the posterior (back) of the nasal cavity is 6.27, the overall range in pH is ∼5.17–8.13 (43). Given that the IC_100_ concentration of niclosamide required to completely prevent viral replication in SARS-CoV-2 infected Calu-3 lung cells as measured by Ko et al (34) as ∼2µM, a 20uM niclosamide solution nasal spray is 10x this IC_100_. If testing in animals and humans reveals 20uM is safe, dose escalation is possible since, according to ***Figure 3B***, the niclosamide concentration can be increased to 30uM at pH 8.07 and 50uM is obtained at only pH 8.31. For the early treatment throat spray, concentrations can be escalated up to 300uM at an orally/buccally tolerated pH of 9.1, which is ∼150x the IC_100_.

Thus, what has been learned is that, when niclosamide is present as a dissolved molecule in aqueous solution, its behavior is predictable by rudimentary concepts of solution chemistry. The theoretical maximum amount in solution is determined by the relative amounts of its pH-dependent unprotonated and protonated forms and follows the Henderson-Hasselbalch and precipitation-pH (pHp) models (***Figures 3***, and ***S4, S5,*** in ***Supplemental Information***). Based on the fitted pKa of 6.87, which is in good agreement with the average of 6.52 from other reported values (11, 57–59), the amount of niclosamide in aqueous solution increases in the higher pH range (of 8-9.5). This is because the fractional amount of the more soluble deprotonated salt form is the dominant species. This pH-dependent behavior provides a mechanism for increasing the bioavailability of niclosamide in simple solutions for more direct application to the mucin-covered epithelia as nasal and throat sprays. Such formulations can be readily prepared simply by dissolving niclosamide powder into buffered aqueous solution, where dissolution times to relative equilibrium take ∼1hr in stirred vials (as in ***Figure 5***). Alternatively, and preferably for ultimate manufacturing, a concentrated ethanolic niclosamide can be mixed by solvent exchange into pH buffers to make the solutions in literally 1-2 seconds, as demonstrated in the rapid injection technique in ***Figure 1***. As pure solutions with no additional solid phase material, these solutions of niclosamide can be made up to 641uM at pH 9.5.

### 5.2. Niclosamide can readily revert to its most stable form, and so reduce its solubility

However, if, and when, precipitated or present as a particulate material, (e.g., micronized or surfactant- or protein- or polymer-stabilized microparticles as observed in our provisional patent work (48)), the amount of niclosamide in aqueous solution is determined by the nature of the crystalline or amorphous material that the local aqueous solution is in equilibrium with, or is moving towards. This is where careful preformulation drug characterization is needed in order to fully understand and predict the nature of niclosamide’s solid forms, its resulting solubility in aqueous media, and hence its bioavailability, in (any) dosage form.

In support of these two major take a-ways, (pH dependent dissolution and reversion to more stable and lower solubility polymorphs) are the studies that explored the apparent solid forms. That is:

- For one commercially available niclosamide powder (AK Sci), after dissolution equilibrium was attained over a period of a few days at pH 9.3, the high concentration solution supernatant was, more often than not, stable despite being in contact with its original excess powder.
- For the Sigma niclosamide, its initially dissolved, high concentration solution was stable for a few hours. However, when left in contact with its original excess powder for longer periods, (e.g., overnight) it converted to a lower solubility solid form.
- For niclosamide precipitated from supersaturation the precipitated particles eventually achieved their own low solubility solid forms and showed a pH-dependent morphology of flat particulate sheets to spiky crystals.
- For the water-precipitate, and acetone and ethanol presumed “cosolvates”, the solubilities were similarly low across the whole pH range.

A summary of this data is now shown in ***Figure 16***. Compared to the original AK Sci material (from ***Figure 3B***) are, the data from ***Figure 8*** (Niclosamide Precipitated from Supersaturated Solution), that from ***Figure 14*** (Sigma Niclosamide after 24 hrs) and the final equilibrium amounts in solution measured for the water, acetone, and ethanol “cosolvates” (***Figure 11***).

**Figure 16.**
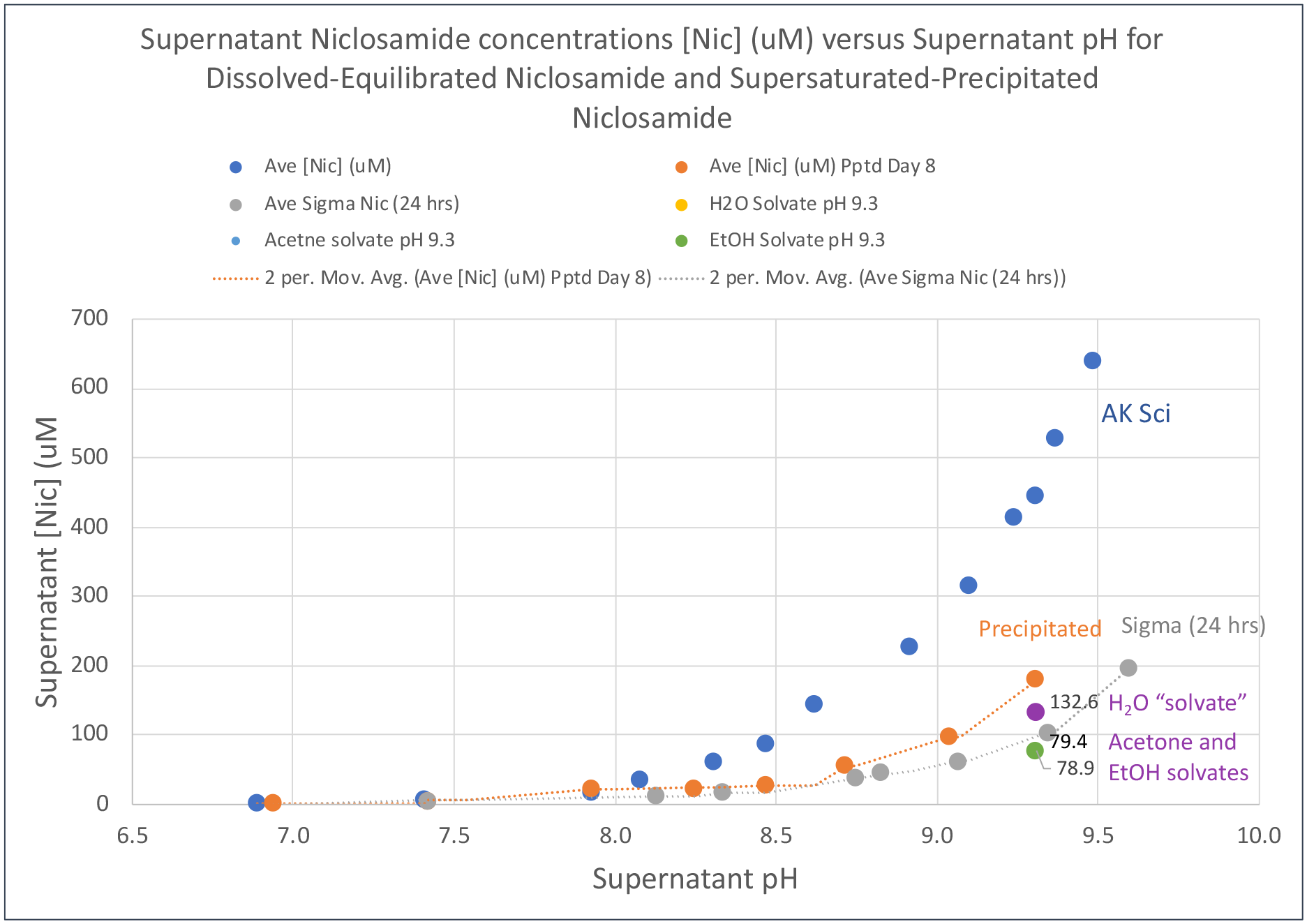
Supernatant Niclosamide concentrations [Nic] (μM) versus Supernatant pH for Dissolved-Equilibrated Niclosamide and Supersaturated-Precipitated Niclosamide plus the water, Acetone and Ethanol (presumed) solvates at pH 9.3. Moving average lines are used to guide the eye.

Thus, as shown in ***Figure 16***, the (probably not surprising) conclusion is that niclosamide can readily revert to its most stable solid forms (--the monohydrates and similar cosolvates identified by van Tonder et al (2, 62)) that have a lower solubility of niclosamide in aqueous solution when in contact with these solid forms.

While still showing the expected pH dependence for solubility i.e., dependent on the acid and salt solution equilibrium, there is a clear distinction and drop in solubility, dependent on the solid form in equilibrium with the species in solution. What this shows is that for both the Sigma niclosamide and the precipitated materials, as well as the presumed “cosolvates”, the amount of niclosamide in actual solution is dependent on the nature of the solid form it is in equilibrium with. Unfortunately for the commercial powders, this information is not available because the manufacturers will not divulge their post synthesis processing to recover the niclosamide material as it is a trade secret.

While beyond the scope of this current study, further studies by Raman and x-ray diffraction FT-IR/Raman, and calorimetry could be carried out and are being planned to gain more insight into the crystal forms of these commercial products and the precipitates and solvates (63).

Thus, when particulate niclosamide is in solution, an equilibrium is set up between the energetics of the solid form (bonding within the crystal form, e.g., crystal lattice energy) and the species in solution. To reiterate, as reported by van Tonder, the most stable forms of niclosamide are the monohydrates that have the lowest solubility compared to anhydrates and other cosolvates. But these forms are not necessarily what are purchased from niclosamide suppliers at AKSci, Sigma or, others that would be of undefined crystal form, and hence solubility, and have not been measured as a function of pH.

As is well known in pharmaceutics, the solubility and permeability behavior of drugs play a major role in bioavailability (76). This is particularly important in local delivery via any nasal and throat spray or nebulized lung inhalant, since, compared to a microparticles suspension, it is only the soluble form that can permeate the mucin layers and reach the epithelia. The value for that solubility is therefore critical for effective drug delivery. In fact, apart from ensuring direct access through the mucin, it was this recognition (of such a low ∼ 2-3uM solubility for niclosamide at neutral pH), that went into the initial thinking (see ***Supplemental Information S1***) and prompted the invention and development of a niclosamide solution as opposed to any microparticle formulation that were all explored in the series of provisional patents (48).

## 6 Summary and Conclusions

With the coronavirus pandemic still raging, more contagious (delta and delta+) variants already in circulation (1), a world-wide average of only 15.4% fully vaccinated (52), and many parts of the world with <1% vaccinated (53), there is still an urgent need for additional mitigation efforts to support current and potential vaccinations. Universal prophylactic nasal and early treatment throat sprays that *“put the virus in lockdown”*, could help prevent infection and reduce viral load in the early stages for unvaccinated populations, as well as vaccinated spreaders where break through infections are now occurring. The niclosamide solutions presented here could represent such universal sprays.

Niclosamide has emerged from multiple drug screens with potential to treat cancer and many other diseases and conditions (5), and now in a broad range of viral infections (6–21). At the cellular level, niclosamide can inhibit three of the six stages of viral infection by: preventing *uncoating* and RNA release from the endosome (19); *inhibiting viral replication* by reducing the amount of ATP available from mitochondria needed for ATP-dependent viral transcription and translation (11); and *interfering with capsid assembly* in the Golgi that then promotes the secretion of non-competent virions (27). The main issue for nose and throat formulations, is how to optimize local bioavailability for a drug that has intrinsic low solubility in aqueous solution (1-2uM) compared to its cellular efficacy (2-3uM) and the fact that the traditional choice for nasal sprays of microparticles cannot penetrate the mucin. It is here that we have focused on what it takes to provide a mucin-penetrating niclosamide solution. The amount of niclosamide in aqueous buffered solution and therefore its bioavailability to infectable and infected epithelial cells, can be increased just by slightly increasing the pH (48).

Experimentation focused on a series of specific aims that measured the pH-dependence of equilibrium supernatant concentrations and dissolution rates of niclosamide in buffered solutions. We compared niclosamide from different suppliers as well as precipitated niclosamide and presumed cosolvates of acetone and ethanol. Nanodrop UV/Vis spectroscopy was used to quantify niclosamide concentrations in supernatant solutions and data was compared to predictions from Henderson Hasselbalch and precipitation pH models. Optimal microscopy was used to observe the morphologies of precipitated and converted niclosamide.

What has been learned is that, when niclosamide is present as a dissolved molecule in aqueous solution, its behavior is predictable by fundamental concepts in physical chemistry and pharmaceutics. Because it has a reported pKa of around 6 to 7 (11, 57–59), and measured in this work to be 6.87, the theoretical maximum amount in solution is determined by the relative amounts of its pH-dependent protonated and unprotonated forms. As shown here, the solubility of niclosamide follows the Henderson-Hasselbalch and precipitation-pH (pHp) models. Basically, the amount of niclosamide in aqueous solution increases with increasing pH, from 1.77uM (and lower depending on the polymorph) at pH 3.66, to 30uM at pH 8. It then showed a more rapid rise in concentration to 90uM at 8.5 and was 300uM at 9.1, reaching 641uM at pH 9.5. This rise in the amount in solution at these more alkaline pHs was because the fractional amount of the more soluble deprotonated salt form is the dominant species. This pH-dependent behavior therefore provides a mechanism for increasing the bioavailability of niclosamide in simple solutions for more direct application as nasal and throat sprays. Such formulations can be readily prepared simply by dissolving niclosamide powder into buffered aqueous solution, or preferably mixed by solvent exchange from concentrated ethanolic niclosamide into pH buffers.

However, if and when precipitated or present as a particulate material, micronized or surfactant- or protein- or polymer-“stabilized” microparticles, the amount of niclosamide in aqueous solution is determined by the nature of the material that the local aqueous solution is in equilibrium with, or is moving towards. This is where careful preformulation drug characterization is needed (see ***Supplemental Information S2***) in order to fully understand and predict the nature of niclosamide’s solid forms, its resulting solubility in aqueous media, and hence its bioavailability in (any) dosage form. Thus, in the preparation of any formulation of niclosamide, especially for local nasal and throat delivery, *niclosamide is not niclosamide is not niclosamide*, i.e., the source and nature of niclosamide can dramatically influence the polymorphism, solubility, amount in solution, bioavailability, and pharmaceutical performance of the drug compound.

Presented here then were preformulation characterizations, conducted experiments, results and discussion that spoke directly to a series of issues in optimizing nasal and throat sprays of niclosamide. They:

- characterized the pH dependent equilibrium and kinetic dissolution behavior of niclosamide from dry powder including niclosamide from various suppliers and co-solvates;
- evaluated its precipitation from supersaturation in aqueous buffered solutions and showed that niclosamide could readily revert to its most stable and lowest solubility hydrated form;
- provided optimized buffered aqueous solutions of niclosamide that are assured to permeate through the protective mucin layers that otherwise protect the airway epithelial cells from microparticles.

Our lead niclosamide formulation is therefore a simple buffered solution that represents a potentially universal prophylactic nasal spray that could stop virus replication at its point of entry, and a higher concentration throat spray, that could reduce viral load as it progresses down the back of the throat. A more detailed presentation and discussion of “Why Niclosamide?”, and its intracellular *virostatic* effects, along with how these measurements are translated into spray products and preclinical and clinical outcomes are discussed in more detail in a white paper in preparation (24). A low dose (20uM) prophylactic solution of niclosamide at a nasally safe and acceptable pH of 7.9 and a (up to 300uM) throat spray at pH 9.1 would be the simplest and potentially most effective formulations from both an efficacy standpoint as well as manufacturing and distribution, with no cold chain. They now just need testing.

## Acknowledgements

Acknowledged are the important discussions with Professors Phil Williams and Jonathan Burley at the School of Pharmacy at the University of Nottingham, UK, and Dr. Carla Gene Rapp, who helped to shape the thinking, interpretation of results, and overall flow of the manuscript. Also, thanks are to Peter Silinski of the Duke Analytical Instrument Facility who analysed niclosamide purity and stability.

## Funding Statement

There was no formal funding from government (e.g., NIH) or other agency. Supplies were paid for from a discretionary account. Regarding salaries, the sole person engaged in all of the work was Professor David Needham PhD, DSc, employed at Duke University in the Pratt School of Engineering in the Department of Mechanical Engineering and Material Science.

## Conflict of Interest Statement

Duke University has assigned the rights for the provisional patent applications to David Needham and his entity, Needham Material Science LLC.

## Author contributions for all authors

David Needham was the sole contributor and sole author

## Data Availability Statement

The datasets generated during and/or analyzed during the current study are available from the corresponding author on reasonable request

## Supplemental Information

Supplemental information contains important aspects of this work that are perhaps outside the scope of the main document, especially regarding space and attention.

A brief description of the initial thinking is given in ***S1***, that went into this particular formulation and some of the initial experiments that led to the “solution”, including: more on the case of Nansonex and Flonase; that Niclosamide’s solubility at physiologic pH is in actually the same range as its efficacy; could Niclosamide be encapsulated or stabilized as nanoparticles?; and given the “solution” to the problem, how much niclosamide is actually sprayed per dose?

As mentioned in the main text, it is important to carry out a preformulation drug characterization focused on the basic molecular properties of the niclosamide, including: its pKa; the solubility of its protonated (S_o_) and deprotonated (S_-ve_) forms; the resulting pH-dependent amount of niclosamide in solution; and its octanol water partition coefficients (LogP, and the pH-dependent Log D) that underly the ability of niclosamide to partition into the various lipid bilayer membranes and exert its proton shunt effect. This is all given now in ***S.2***

Additional supplemental information is also provided in ***S3*** for the Calibration of Standards, that is important to show because the UV/VIS spectra of niclosamide in aqueous solution at pH 9.3 are actually new.

### S1. Initial thinking and early experimentation

For completion, this section briefly describes some initial thinking and early experimentation that generated a series of provisional patent applications (2) and guided the development of the simplest and most optimized formulation, the pH buffered solutions.

#### S1.1 Niclosamide readily precipitates as a visible precipitate

In the very first and simplest experiment, to evaluate the potential for niclosamide to make solutions or precipitate, ethanolic niclosamide solution was injected into excess aqueous (anti-solvent) buffer by using the same solvent exchange technique we have used previously for making micro and nanoparticles (3), (see 3.2 Methods, 3.2.1). As shown in ***Figure S1***, at physiologic pH, at a final concentration of 1mM niclosamide precipitates, and forms large, and aggregated microparticles that present as a “fluffy”, gel-like, precipitate, that is visible by eye (1) (see main text, microparticle images at pH 7 in ***Figure 9***). Such precipitate can be seen all the way down to a few micromolar concentrations of niclosamide. The mucosa is designed to protect the nasal epithelial cells and prevent the permeation of particles with a size greater than a few 100 nanometers (4–6). Only small nanoparticles, micelles, and dissolved molecules of niclosamide in solution can get through.

**Figure S1.**
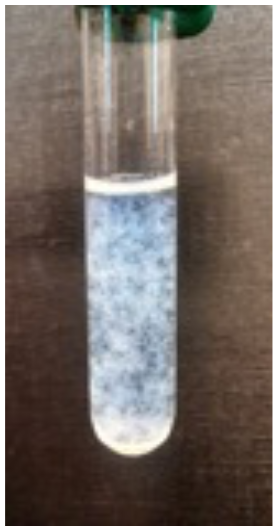
1mM Precipitated Niclosamide forms a visible “fluffy” precipitate, especially when shaken, that would be unsuitable for a nasal spray (1)

In the initial thinking for formulation design for this nasal and buccal epithelial application, it was therefore feared that, while easy to make, particulate suspensions of niclosamide would simply be trapped at the top surface of the protective mucin layer and so would not be fully optimized for a nasal or throat spray. That is, microparticles that cannot penetrate the mucin barrier can only provide the small fraction of the low solubility niclosamide (1μM – 2μM) in solution and so do not optimally deliver molecular-niclosamide to the underlying epithelial cells. However, microparticle formulations such as Nasonex and Flonase do exist for nasal applications, and so what is the situation here?

#### S1.2 The case of Nansonex and Flonase that could inspire microparticle formulations

While the microparticle-micronized drug concept for drug delivery is clearly a well-established option, especially for nasal sprays, is it really optimized for niclosamide? It is here that considerations of crystal morphology and their pH dependent solubility are key determinants in the design and ultimate performance-in-service. As an example, in stark contrast to the 1-2μM solubility of niclosamide and its 2-3μM IC_100_ virostatic efficacy (7, 8) micronization does work for similarly low solubility drugs like mometasone furoate and fluticasone propionate i.e., as Nansonex (9–11) and Flonase (12–14). Consider first the formulations themselves and the amount of micronized and soluble drug present compared to their efficacy. For mometasone furoate, according to the FDA IND approval (10), Nasonex Nasal Spray in aqueous medium, contains: 50 micrograms of the API mometasone furoate; Glycerin, microcrystalline cellulose and carboxymethylcellulose sodium, sodium citrate, citric acid (to control pH) benzalkonium chloride (preservative), and polysorbate 80. It has a pH between 4.3 and 4.9. Mechanistically, Mometasone furoate diffuses across cell membranes to activate pathways responsible for reducing inflammation. Each actuation of Nasonex delivers 50 mcg of mometasone furoate in 100uL of formulation through the nasal adapter. Thus, the mometasone furoate concentration per spray (50 x 10^-6^g/ 100 x 10^-6^L) x 1/521.43 g/mol) = 958μM, i.e., almost 1 mM. As micronized material this is in presumable equilibrium with its aqueous solubility of 20.7μM (calculated from chemaxon).

By comparison, the relative trans-activation potency (EC_50_) for mometasone furoate is 0.069 ± 0.021 nmol/L, i.e., the amount required to induce GR-mediated transcription of a synthetic target gene regulated by a glucocorticoid response element is delivered at 14 million times more drug than its EC_50_. Similarly for fluticasone propionate with an aqueous solubility of 22.8µM, its EC_50_ activity is 0.320 ± 0.04 nmol/L and so Flonase delivers 3 million times more drug than needed for activity. As is clear, most of this drug is in the form of micronized mometasone furoate or fluticasone propionate, that cannot get through the mucin that covers the underlying epithelia. And so, the key mechanism here for drug transport is simply the diffusion of the soluble fraction, (21μM or 23µM) of drug that can permeate through the mucin. Thus, while Nasonex and Flonase might provide good examples and motivation for micronized drug formulations for nasal application, they only work because of the huge 30,000 to 6,000 times greater bioavailable aqueous concentration of the drugs compared to their nanomolar efficacy.

Thus for these medications, it’s the amount of drug in solution at 20,000 times the effective dose at the receptor that gets to the underlying cells to have its corticosteroid receptor action, while the several million times excess micronized drug just sits on the mucus that is being replaced every 15 minutes (15) and so is acting as a possible depot until it is excreted.

#### S1.3 Unfortunately, Niclosamide’s solubility at physiologic pH is in the same range as its efficacy

Since niclosamide is such a low solubility drug at physiologic pH, (1μM – 2μM) and its anti-viral efficacy was determined to also be in this same range of 1µM (16) in Vero 6 cells and 2-3 times less in the more appropriate lung Calu-3 cells (∼ 2-3µM) (7, 8), a pH 7 aqueous solution of niclosamide at equilibrium with any particulate niclosamide may not be sufficient for optimal efficacy. It really does not matter how much microparticles of niclosamide are sprayed intranasally, the bioavailability is essentially set by the solubility of the niclosamide solid material in the formulation at the environmental pH. Given that nasal mucosa has a refresh rate of ∼21 mins (15), even as a depot of material, microparticles would likely not dissolve sufficiently well or quickly enough in the limited amount of water present in the mucosa to provide optimal niclosamide to the underlying epithelia. Thus, while microparticle niclosamide may provide some efficacy because it does provide some but limited amounts of niclosamide in solution, it is clearly not an optimally bioavailable formulation.

#### S1.4 Could Niclosamide be encapsulated or stabilized as nanoparticles?

Since March 2020, efforts were focused on ways to increase the amount of niclosamide in a sprayable suspension stabilized by a series of common surfactants, polymers, and preservatives routinely used in, for example, mouthwash, nasal sprays, and eye drops (2). The goal was to make nanoparticles that could permeate the mucin, because, with a particulate cut off of ∼0.5 um (6) microparticles of ∼1-10 um diameter would be unlikely to provide optimal delivery of niclosamide.

Attempts were therefore explored to stabilize the large visible particles, seen in ***Figure S1***, into something smaller that could stabilize niclosamide nanoparticles and so could in principle permeate the mucin. We have previously used a solvent exchange technique to introduce ethanolic niclosamide solutions into excess aqueous anti-solvent to form drug delivery particles for cancer (3, 17). Using this technique for niclosamide and a range of surfactant, polymer and protein stabilizers, many well-stabilized micro- and even nano-particle suspensions were prepared. A series of provisional patent applications were obtained (2). However, while of low aqueous solubility, niclosamide is still soluble enough for any precipitated nanoparticles to ripen and become larger microparticles, which, again, would merely stick to the top of the mucin layer and so be unsuitable (or at least not optimized) for nasal sprays. Also, as presented in Results (4.3, 4.4, and 4.5), and recognized and characterized by others (18–21) *“niclosamide is not niclosamide is not niclosamide”.* That is, commercially available niclosamide that may be initially purchased and used in a formulation, but it is not necessarily the most stable and therefore least soluble form, can convert in aqueous media to low solubility material over time. As well-recognized in pharmaceutical preparations (22), what this means, formulation-wise, is that, even if a microparticle formulation is made, if resuspended in aqueous buffer, it could readily convert to a low solubility morphology and so become less bioavailable in solution,

Examining individual surfactants in the commercial mouthwash and hydrating nasal sprays that are included as stabilizers gave micellized niclosamide and somewhat stabilized microcrystals. The quaternary ammonium preservatives also stabilized positively charged ionic-complexed nanoparticles (of ∼0.2um - 0.5um in diameter). These latter particles could, in principle, adhere to the negatively charged mucus. However, they were found to be so stable that they would not sufficiently dissolve. As for micellar solutions (Tweens Poloxamers), while readily prepared, such relatively high surfactant concentrations could be toxic to the nasal epithelium.

Again, this is not to say that the limited amount of niclosamide in equilibrium solution with a microparticle suspension stuck at the mucin surface would not reach the underlying epithelial cells to some extent and show some efficacy, (as it has been encouragingly shown to do with micronized niclosamide (23)). It is just that this could be optimized by using a much simpler solution at higher but still tissue-tolerated pH. In contrast, 20uM to 300uM niclosamide solutions (with no microparticles) are expected readily diffuse through and permeate the mucin to more-optimally deliver greater amounts of niclosamide to the underlying epithelial cells, as presented next.

#### S1.5 A “Solution” to the Problem: How much niclosamide is actually sprayed per dose?

As described in S2. Preformulation drug characterization, and demonstrated in the results section (4.1), an evaluation of the physicochemical properties of niclosamide revealed a much simpler solution (literally); one that could be obtained by slightly increasing the pH (to pH 8) of the aqueous media that niclosamide was dissolved into. This provides niclosamide concentrations of 20uM - 30uM that are 10x the efficacious virostatic levels of 2uM-3uM (8, 16) in Calu-3 cells, and that we have shown effectively reduce the ATP produced in host airway epithelial cells at levels that are also non-toxic to these cells (24). Therefore, prophylactic nasal spray solutions of 20uM - 30uM are expected to be safely administrable at pH 8, which is within the normal nasal pH range. Niclosamide solutions of up to 300uM niclosamide have been obtained at pH 9.1, which is on the same order as the pH of green tea or commercially sold alkaline water and so for oral administration this pH is also expected to be safe, and so could potentially be used as an early treatment throat spray.

For the treatment of worms, 4 x 500mg of Yomesan tablets are thoroughly chewed in the mouth or mixed to a paste in 30mls of tap water and swallowed. In comparison to these 2 grams of niclosamide that the buccal and throat epithelia is exposed to during Yomesan administration, for the throat spray, a single spray of 100uL of 300uM niclosamide is only 9.8 micrograms per spray, i.e., 204,000 times less niclosamide. In reality, it is not this dramatic a difference because, as is one of the main themes of this paper, we have to take into account what is actually *bioavailable* to the buccal epithelium as molecular dissolved niclosamide and not just particulate niclosamide tablet. So, what would this represent? A simple calculation provides the answer.

If the pH of tap water (and saliva) is ∼ pH 6-7, then, as measured in the Results section, the amount of niclosamide in solution at this pH is ∼2uM. Therefore, the amount of niclosamide that could potentially dissolve out of the 2 grams of tablets in 30mLs at 2uM niclosamide, is 60 nanomoles or 19.6 micrograms (Mwt niclosamide is 327.1 g/mol). Compare this now to the 100uL of 300uM throat spray dose of 9.8 micrograms, which is similar, but still 50% less than the approved oral tablets. So why not just chew the Yomesan tablet? Because it is 204,000 times more niclosamide than you need, and, if swallowed, does have GI side effects that would make it somewhat intolerable on a regular basis and so not that patient compliant as a prophylactic or early treatment regimen. In fact, this is the basis for the demonstration to be reported in a separate note (25) that niclosamide can be readily extracted from existing approved commercial Yomesan (Bayer) tablets to higher concentrations than in tap water or saliva if the pH is raised.

For the prophylactic nasal spray at 20uM niclosamide, 100uL of 20uM niclosamide solution is only 0.65ug per spray, i.e., 3 million times less in total mass and still ∼32x less in terms of bioavailable niclosamide. These simple calculations actually emphasize the main theme of this paper, that, solubility is everything when it comes to epithelial bioavailability.

### S2. Preformulation Drug Characterization (pK*_a_*, S*_w_*, LogP)

This section provides a brief theoretical and rudimentary basis that lays the foundation for the niclosamide solution formulation. It is focused on the basic molecular properties of the compound, including its pKa, the solubility of its protonated (S_o_) and deprotonated (S_-ve_) forms, the resulting pH-dependent amount of niclosamide in solution, and its octanol water partition coefficients (LogP, and the pH-dependent Log D) that underly the ability of niclosamide to partition into the various lipid bilayer membranes and exert its proton shunt effect. It is here that a careful preformulation drug characterization is needed in order to fully understand and predict the nature of niclosamide’s morphological forms, its resulting solubility and concentrations that can be attained in aqueous media, especially as a function of pH, and hence its bioavailability in (any) dosage form.

As shown in ***Figure S2***, Niclosamide (5-Chloro-N-(2-chloro-4-nitrophenyl)-2-hydroxy-benzamide), is a chlorinated salicylanilide, that contains a zwitterionic nitro group and an ionizable OH. The most important moiety for the current application is the ionizable OH, and so analysis starts with the effect of deprotonation of this salicylic OH (Nic_OH_). Deprotonation gives the negatively charged salt (Nic_-ve_), which occurs over a certain range of pH and is determined by the molecule’s pKa. The reason this is important is that, with a pKa close to neutral pH, the protonated acid is the most prevalent species at any lower pH, such as in the nasal cavity, unless buffered with higher pH solution. The Nic_OH_ acid species is expected to have a much lower solubility than the charged salt and so determines the amount of niclosamide in solution. Therefore, by measuring the solubility of the acid and determining the amount of niclosamide in solution we can evaluate this pH-dependent behavior and obtain the best fit pKa. It is therefore this weak acid pKa that enables the new higher concentrations of niclosamide to be prepared in simple buffer solutions. The questions answered in this study then are: *“How much does the amount of niclosamide in solution increase with increasing pH?” “How high does the pH have to be to achieve niclosamide concentrations that are well in excess of the therapeutic levels of 1µM – 2µM?”* and, eventually, *“Are these levels expected to be safe in the nasal and buccal/throat epithelium?”*

**Figure S2.**
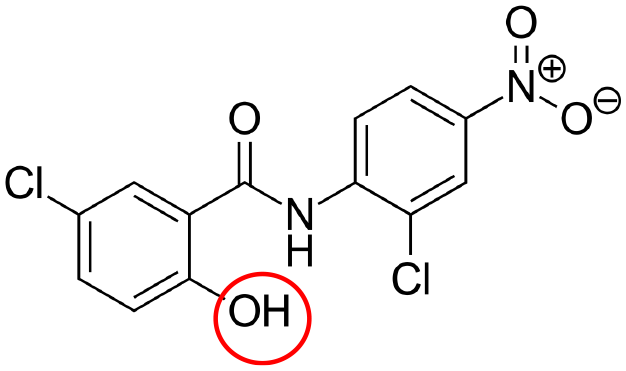
Niclosamide (5-Chloro-N-(2-chloro-4-nitrophenyl)-2-hydroxybenzamide) contains an ionizable OH and a zwitterionic nitro group. It is the ionizable group that provides for the much higher aqueous solubility as a function of pH.

Taking the basic molecular properties in turn provides rudimentary and important quantitation of the properties of this drug that are the basis for the new and simple solution formulation and spray, its mechanism of action, and that should really be considered in any other nano or microparticle formulation of niclosamide.

#### S2.1. pK_a_

The pKa is defined as the negative base-10 logarithm of the acid dissociation constant (Ka) of a solution). It is an important parameter to consider here because pKa and pH are equal when half of the acid has dissociated. Although experimentally challenging for such a low solubility molecule as niclosamide, the pKa can be measured experimentally (26) or calculated theoretically (27); both are available in the literature, and a range of values are reported for niclosamide. Jurgeit et al (*28*) report an estimated pKa of ***5.60*** as given in (29), calculated using solubility data, also confirmed at U.S. EPA ECOTOX database (30) related to Pesticide Risk Assessment. Drugbank.ca, quotes the pKa as ***6.89*** and references calculations by Chem Axon. Similarly, at ACD/iLabs, niclosamide is calculated as having a pK_a_ of ***7.2***, and in (31) the phenolic moiety has quoted as having a pKa = ***6.38*** (32). Thus, the literature has a range of reported pK_a_ values from ***5.6 to 7.2***, and the average of all thesevalues is ***6.52***.

Once the pKa is measured (or estimated) the well-known Henderson-Hasselbalch equation calculates the fractions of each species, i.e., the protonated acid (Nic_OH_) and the deprotonated salt (Nic_eve_). For a weak acid like niclosamide this is simply:

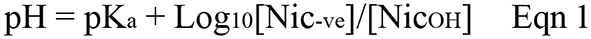

This equation can be represented as:

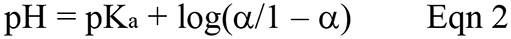

where *α*is the fraction ionized species as the deprotonated salt, Nic_-ve_. As outlined in (33) the fraction ionized can then be obtained as:

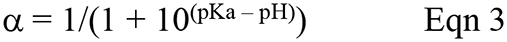

As shown in ***Figure S3***, for a known pKa, the Henderson-Hasselbalch equation allows a plot to be made of the expected fractions of each species: *α* is Nic*_-ve_* (orange symbols) and 1-*α* is Nic*_OH_*, (blue symbols), exemplified for pKa of 6.87 (derived from experiment, see later, Results, 4.2 Equilibrium Dissolution of Niclosamide versus pH).

**Figure S3.**
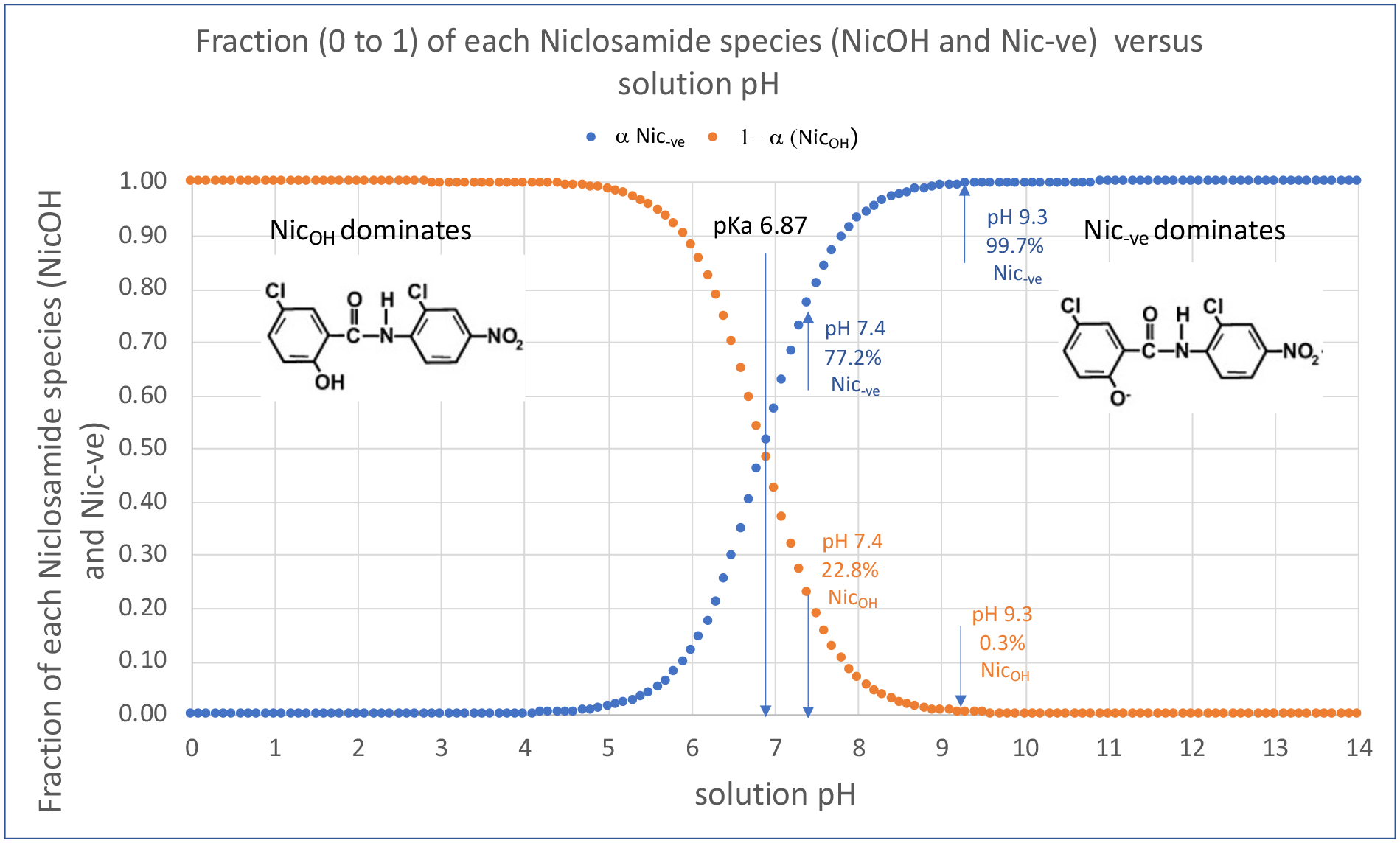
The Henderson-Hasselbalch equation calculates the fractions of each species as a function of pH, shown here for a nominal pKa (derived from experiment) of 6.87.

As can be seen in ***Figure S3*** at low pH, the dominant species is the protonated acid Nic*_OH_*. Then, with increasing pH, Nic_OH_ deprotonates and the salt Nic*_–ve_* starts to become more prevalent. As seen in ***Figure 3A*** and ***B***, in the main paper, fitting the pHp data for a measured protonated acid solubility, S*_o_* of 1.77uM, niclosamide gave a pKa of 6.87; so this, in and of itself, is a way to “measure” pKa. Thus, if this value is used as an example, there is 50% of each niclosamide species present in solution at pH 6.87. As can be seen in ***Figure S3***, at low pH, (less than pH 4), the low solubility acid represents 100% of niclosamide in solution, and the percentage amount of this species is expected to decrease with increasing pH. Similarly, the percentage of the more soluble charged salt is expected to increase with increasing pH; it becomes the 99% dominant species at pH 8.9 and 100% dominant species at ∼pH 10.

Thus, the key result from this graph for the new application of niclosamide as a nasal and throat spray solution, is that a niclosamide solution concentration equilibrium is expected between the acid and the salt for any solution pH between pH 4 and 10. For example, at physiological pH of 7.4, the acid and salt are 24.4% and 75.6% of the niclosamide in solution, respectively. At pH 9.3, Nic*_-ve_* is 99.7% of the acid salt pair and the low solubility acid is only 0.3%. Note: it is the high solubility salt (as a lipophilic anion (28)) that does the proton shunting across membranes.

Overall, then, the amount of niclosamide in aqueous buffered solution is expected to be higher at higher pH. This is the basis for the current solution formulation of niclosamide. As described later, there are limits as to the range of pH that can be applied in the nasal-, but less so in the oral-pharynx. However, at first glance, this pKa of 6.87 does appear to give a workable range and a potentially useful formulation for a niclosamide solution.

#### S2.2. Aqueous Solubilities and amounts of niclosamide in solution

The next challenge is to obtain the maximum concentrations of niclosamide in solution as a function of pH. For this we need to measure or estimate the solubilities of each species, or at least one of them, i.e., the low solubility acid (S_o_) as later, in Eqn 4. Obviously, the reason niclosamide solubility is so important is because it determines the solution concentration, and hence bioavailability, of niclosamide that can permeate through the mucin to the all-important layer of epithelial cells.

Values of aqueous solubility of niclosamide in the literature are again widely distributed. One reason may be the different polymorphic forms that are tested, as well as pH (that is often not specified). As quoted in a paper by Graebing (34), the solubility of niclosamide in water is 5-8 mg/L (14) which is, a relatively high, 15.5uM to 24.5uM. In contrast, an EPA document says: *“Niclosamide is practically insoluble in water”* and gives a value of 1.05 x 10^-5^ g/100 mL which is 0.321uM. As characterized for niclosamide in a series of papers by de Villiers et al (18–21), polymorphism, hydrate versus anhydrate, and cosolvates can change the solubility, dissolution, and chemical stability of the drug compound. These are probably the most reliable values (21) where the solubility of the crystal forms was measured in distilled water and found to decrease in the order: anhydrate ⪢ monohydrate HA > monohydrate HB with the monohydrate being the most stable form. The values reported were: Anhydrate 13.32 ± 3.18μg/mL; Monohydrates HA 0.95 ± 0.06 μg/mL; Monohydrates HB 0.61 ± 0.09 μg/mL. These values convert to 40.4uM, 2.9uM and 1.87uM, respectively.

As described here in the Experimental section in Methods and Results, it has been possible to obtain some estimate of the limiting solubility of niclosamide from nanodrop UV/Vis by evaluating two noisy scans, i.e., one for the blank buffer and one for the niclosamide solution at pH 3.66, where the niclosamide solubility is expected to be its absolute lowest. Again, this solubility (for the AK Sci niclosamide) was measured to be only 1.77uM ± 1uM, and so is actually in good agreement with van Tonder and de Villiers (21). Even though the pH was not measured or reported in van Tonder’s paper, a deionized water solution of niclosamide has a pH of ∼6.5, (as measured in the present study) and the solubility obtained at pH 6.89 of 3.8uM ± 1uM, is in good agreement.

Lastly, there appears to be only one mention of the pH-dependent solubility of Niclosamide in the literature. The Pesticide Handbook (35) reports that the solubility of Niclosamide at pH 6.4 as 1.6mg/L (which is 4.89uM) and at pH 9.1 is 110mg/L (which is 336.3uM), suggesting that the solubility of niclosamide can vary by at least a factor of almost 100x depending on pH between 6.4 and 9.1. This data supports the new findings here now measured across the whole pH range. In any event, care must be taken to specify pH when reporting any solubilities for niclosamide.

#### S2.3. The precipitation pH (pHp)

These two properties of pKa and water solubility are brought together by combining them with Henderson Hasselbalch analysis to give the precipitation pH that allows us to determine the expected amount of niclosamide in solution across the whole pH range. The precipitation pH (pHp) is the pH below which an acid (or above which a base) will begin to precipitate (with all the necessary caveats associated with stochastic homogeneous and/or heterogeneous precipitation from supersaturation). It is related to the pKa of the drug and its concentrations by the equation:

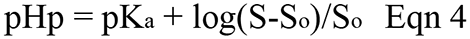

where, S_o_ = the molar solubility of the undissociated acid (Nic*_OH_*), and S = the molar concentration of the salt form in solution (Nic_-ve_). The pHp therefore provides a precipitation limit for the amount of niclosamide in solution as a function of pH, as we might want for optimized nasal and throat sprays. ***Figure S4*** shows a plot of this limit to niclosamide solubility (as pHp) versus Niclosamide concentration for (***A***) a range of pKas from 5.5 to 7.0 for a constant S_o_ of 1µM; and (***B***) a series of solubilities of Nic_OH_ (S_o_) of from 0.25μM to 5 µM and a constant pKa of 6.87.

**Figure S4.**
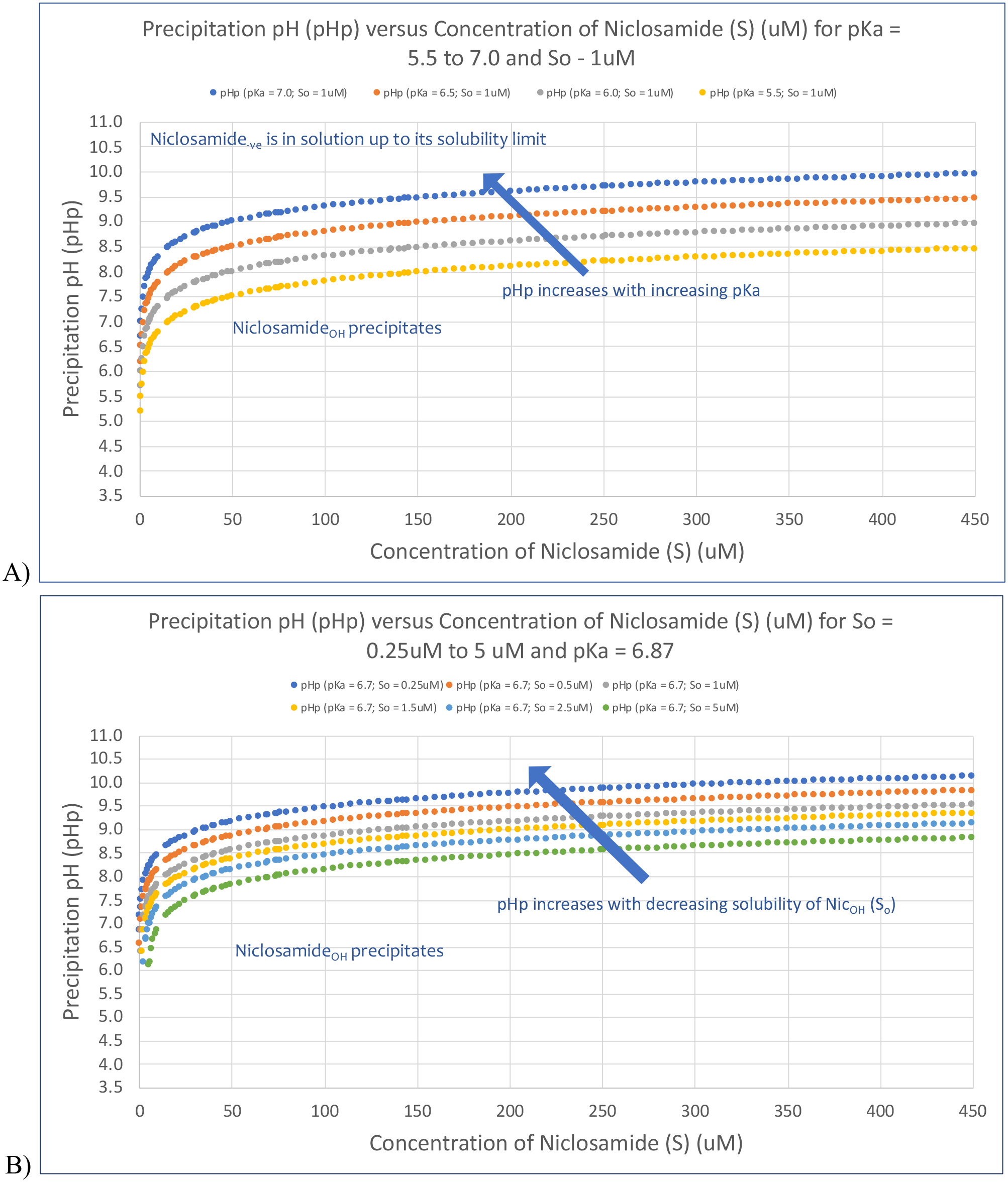
The effects of two parameters: Precipitation pH and pKa of niclosamide on its solubility limits. A) Precipitation pH versus Concentration of Niclosamide (mM) as a function of pKa (5.5-7.0) and constant nominal solubility of Nic_OH_, S_o_ of 1uM; B). Precipitation pH versus concentration of Niclosamide (uM) as a function of solubility of Nic_OH_, S_o_, (0.25uM – 5.0uM) and constant nominal pKa of 6.87

What this graph shows is that the pHp provides a precipitation limit for the amount of niclosamide in solution as we might want for a throat or nasal spray. As indicated in ***Figure S4A***, in the region of the graph above the lines, Niclosamide is in solution, predominantly as Nic_-ve_ the negatively charged salt (up to its solubility limit, measured later to be ∼640μM at pH 9.5); below the lines, niclosamide precipitates. Thus, with the two niclosamide species, there are lower and upper limits to solubility. Niclosamide has such a limited amount in solution. Of course, the precipitation, when initiated from super saturated solution, will depend on the degree of super saturation and time allowed for the precipitation to happen if in this part of the plot. Such supersaturations and the nature of the precipitated material are therefore investigated (Methods, 3.2.5 and 4. Results, 4.4).

As we will see in the results of later experiments, with S_o_ measured to be ∼1.77μM and with a value for the pKa of 6.87, we obtain a very good fit to the experimental data. The overall result though is that for a given niclosamide acid solubility, the pHp increases with increasing value for the pKa. Similarly, as shown in ***Figure S4B***, a plot of pHp versus Niclosamide concentration for a range of Nic_OH_ solubilities (S_o_) (0.25μM to 5μM) and constant pKa of 6.87, shows that the pHp increases with decreasing solubility of Nic_OH_. As expected, it is S_o_ that affects the position of the curve and does so to a similar extent as the range of possible pKas. Clearly, the accuracy of the concentration measurement at these very low solubilities is quite critical.

Basically, the curve can be shifted to higher pHps by either decreasing the value of the solubility or increasing the value of the pKa. Thus, to what extent we can fit theory to the data depends on the accuracy of not only the concentrations as a function of pH, but also on the accuracy of the measured solubility of the very low solubility Nic_OH_ species and, if available, on measuring or calculating the pKa itself.

#### S2.4 Comparison between HH curves, pHp, and amount of niclosamide in solution

To complete this theoretical assessment, plotted in ***Figure S5*** (red symbols) is the pHp curve, giving the maximum amount of niclosamide expected to be in solution (uM) versus the solution pH (right axis) using the values derived from experiment of pKa = 6.87, S_o_ = 1.77uM.

**Figure S5.**
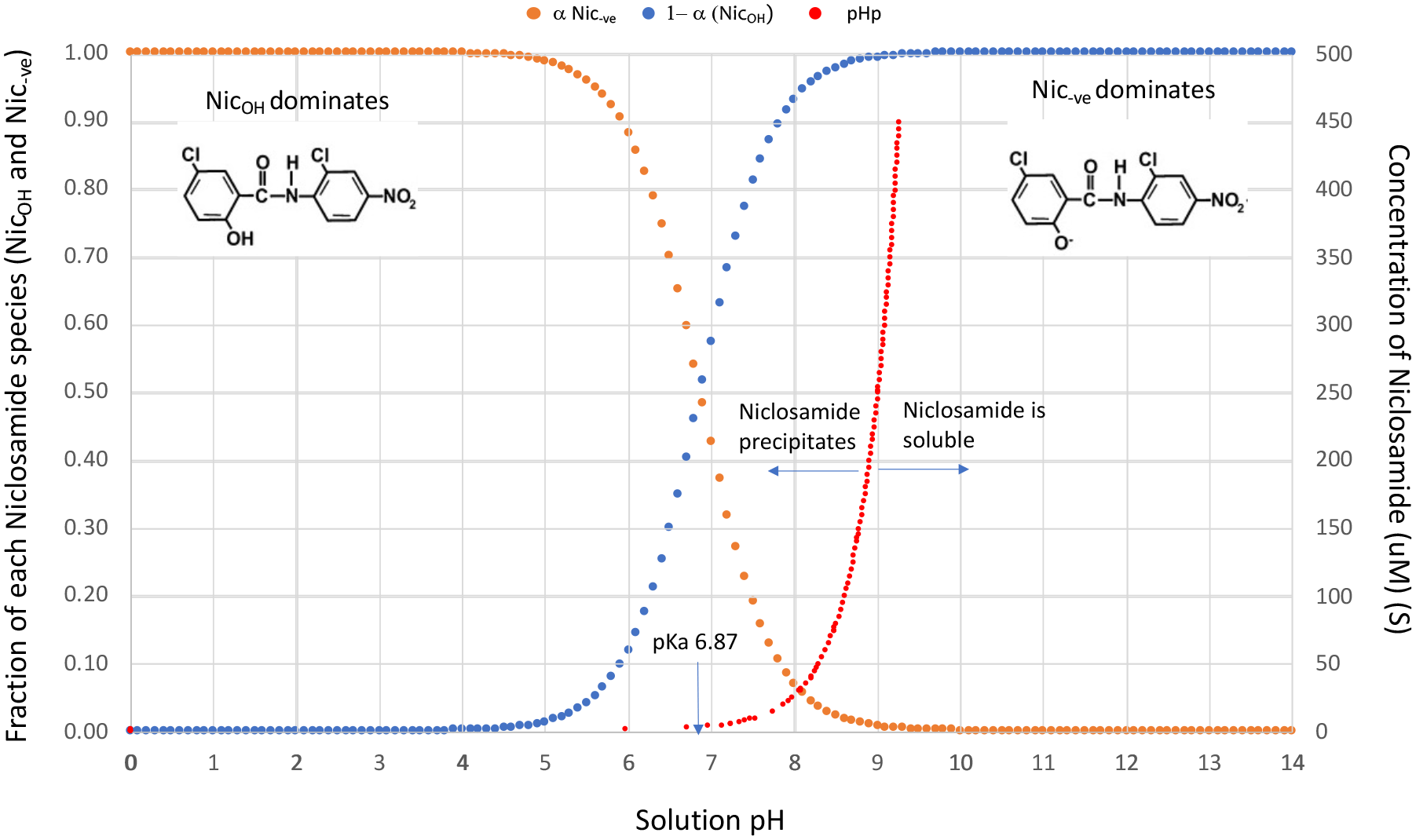
Comparison between Henderson Hasselbalch (HH) curves (orange and blue for the Nic_OH_ and Nic_-ve_ species) (left axis), and the precipitation pH (pHp) (red symbols) (right axis) for niclosamide as a function of solution pH for pK_a_ = 6.78, S_o_ = 1.77uM.

It is superimposed on the fractions of each Niclosamide species (Nic*_OH_* and Nic*_-ve_*) (left axis) from the Henderson Hasselbalch analysis (from ***Figure S3***).

What this graph demonstrates is that the amount of niclosamide in solution (as measured below in the supernatants of dissolving material by UV/Vis) only starts to increase significantly beyond the pKa of 6.87. This analysis implies that, at pH 6.87 the 50% of the low solubility acid determines the total amount of niclosamide in solution, i.e., from Eqn 4, the other 50% of niclosamide is the highly soluble deprotonated salt that matches the amount of the low solubility acid in solution. Thus, the expected amount in solution is 2 x 1.77uM = 3.54uM. Below the solubility limit of Nic_-ve_ and above the solubility limit of Nic*_OH_*, the amount of Nic*_OH_* in solution is only 1.77uM, and the Nic*_-ve_* makes up the rest. Therefore, as shown in ***Figure S5***, for a given pKa of niclosamide of 6.87, the position of this curve, that delineates the amount of niclosamide in solution across the whole pH range, is determined by the value for NIC*_OH_*, the low solubility acid, which, again, is measured here to be 1.77uM.

#### S2.5. LogP

Finally, the most important parameter with respect to mechanism of action of niclosamide on cellular membranes is its LogD, which gives the extent niclosamide partitions into octanol versus water as a function of pH. This parameter, along with its size as molar volume (V_m_ in cm^3^/mol) and its topological polar surface area (TPSA in Å^2^), determines niclosamide’s propensity to partition into lipid bilayer membranes, actually as the parameter, LogB (LogBilayer). Gobas et al (36) compared LogP and LogB for a series of 27 selected halogenated aromatic-hydrocarbons measured in dimyristoylphosphatidylcholine (DMPC) bilayers. Their data showed that, while the octanol-water partition coefficient (LogP) increased linearly with molar volume, the membrane-water partition coefficients (LogB), increased initially with LogP for the smaller molecules, but then followed a more parabolic relationship with respect to larger V_m_. The maximum LogB occurred for solutes with molar volumes of ∼300 cm^3^/mol, after which, their ability to partition into bilayers went down; larger molecules could simply not fit into the liquid-crystalline bilayer structure of the lipid acyl chains and are therefore excluded, as we demonstrated for long chain alkanes and squalene in 1983 using black lipid films, (37).

For the case of such a small molecule as niclosamide, (V_m_ = 202.5±3.0 cm^3^, ACD/iLabs) and a low topological polar surface area of 95.15Å^2^ (Drugbank), (where any value below 140Å^2^ is considered hydrophobic enough to partition into the membrane interior), it is well within the parameters for membrane partitioning. Thus, since we are focused on its pH dependence, it is now the LogD that is an indication of its ability to partition into lipid bilayer membranes and exert its proton shunt effect. Normally, it is the extent to which the molecule is charged or not, since charges are not well partitioned into low dielectric (dc) media like the inside of lipid membranes where the dc is 2. However, it is here that niclosamide has its advantageous property, to delocalize the charge and become a lipophilic anion (28). As given at ACD/iLabs, LogD niclosamide (for their calculated pKa of 6.89) is 4.55 at pH 0.01 and at pH 7.2 it is still 4.23. At pHs well above its pKa, the LogD is still 3.5. The reason its LogD does not decrease appreciably after it is deprotonated is, again, because the negative charge is delocalized within the molecule in internal hydrogen bonding, such that it becomes a lipophilic anion and so can act as the all-important intramembrane proton shunt (28).

### S3. Calibration Standards for the amount of Niclosamide in solution versus pH

#### S3.1 Experimental

Calibration standards (of 20uM, 50uM, 100uM, 150uM, 200uM, 250uM and 300uM) for the UV/Vis experiment were prepared by injecting 100uL of each ethanolic niclosamide solution into 10 mLs of the 9.3 pH buffer giving a 1:99 dilution and a final 1% ethanol solution of niclosamide. To achieve each niclosamide concentration, the 30mM stock solution was made by weighing 100.09mg of AK Sci niclosamide powder (i.e., 98.13mg of niclosamide, allowing for the ∼2% impurity that is filtered out later) and dissolving in ethanol up to 10mLs total volume. Subsequent dilutions of 2mM, 5mM, 10mM, 15mM, 20mM, and 25mM ethanolic niclosamide were made in 2mL final volumes, and 100uLs of each ethanolic niclosamide solution were injected into individual vials of 10mLs of pH 9.3 buffer. Again, this meant that there was a residual 1% ethanol in each standard. The Yalkowski model for solute solubility as a function of cosolvent-water mixtures (in this case ethanol) (38–40) shows that 1% ethanol would only increase the solubility of niclosamide by ∼4% compared to pure buffer. Actually, compared to a simple weighing of powder directly into aqueous buffer (20uM is only 0.066mg/10mLs)), this technique provides a very convenient way to make more pure solutions. Also, the small amount of insoluble impurity that appears to be in the AK Sci product, and does not dissolve in ethanol, and can be filtered out prior to making the solution using a 0.22um filter.

A calibration for niclosamide concentration (y) versus UV absorbance at 333nm (x) was first obtained on a Thermo Scientific Nanodrop 1000 UV/Vis Spectrophotometer over the range 25uM-300uM niclosamide, providing a linear fit of y = 0.0014x - 0.0006. It was used to calculate the concentrationsof niclosamide from the sample absorbance. Weighing such small amounts to create 10mLs or even 20 mLs of sample for the standards was difficult and even beyond the sensitivity of the mg balance (10mLs of a 100uM Niclosamide solution requires weighing 0.327mgs). Rather, solutions were made by a solvent injection technique (see main text 3.2.1) and so gave much more accurate solution concentrations.

#### S3.2 Results

Calibration standards were made up in 10mL volumes in 20mL scintillation vials at nominally 20uM, 50uM, 100uM, 150uM, 200uM 250uM and 300uM concentrations. ***Figure S6*** is a photographic image of the series of calibration standards made by solvent exchange technique in pH 9.3 Trizma buffer.

**Figure S6.**
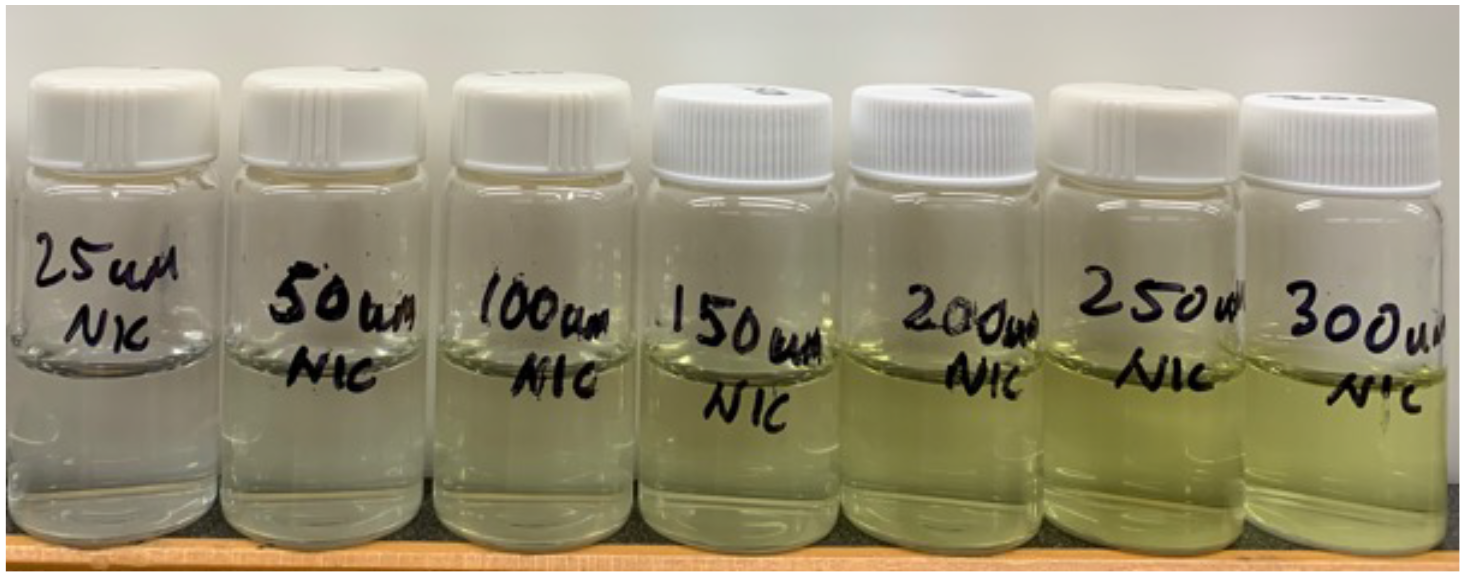
Photographic image of the series of calibration standards as 10mL samples showing the increasing yellowness of the samples with increasing niclosamide concentration from 25uM to 300uM all

It shows, quite visually, how the concentration of niclosamide creates a greater yellowness with increasing pH of the solution. Because niclosamide is usually thought to be relatively insoluble in aqueous media, calibration spectra are usually obtained in methanol, methanolic 0.1 N HCl, and methanolic 0.1 N NaOH (41). The dominant peaks in methanol are at 333nm and 377nm. While niclosamide in methanolic 0.1 M NaOH continues to show the double peaks, in methanolic 0.1 M HCl, where the salt is lost and the prevalent species is the protonated acid, there is only a dominant 333nm peak. The constant peak at 333nm was therefore chosen to report on niclosamide concentration in aqueous solution. Here, we introduce for the first-time spectra for niclosamide at high pH but still just in aqueous media.

UV/Vis Absorption spectra are shown in ***Figure S7***, from which the average absorption values were obtained at 333nm and 377nm. As shown in ***Figure S7***, the niclosamide UV/Vis spectra from 25uM to 300uM all show the characteristic double peak profile at this pH of 9.3, with *λ*_max_ maxima at 333nm and 377nm, although the maximum in aqueous pH 9.3 buffer was closer to 335nm.

**Figure S7.**
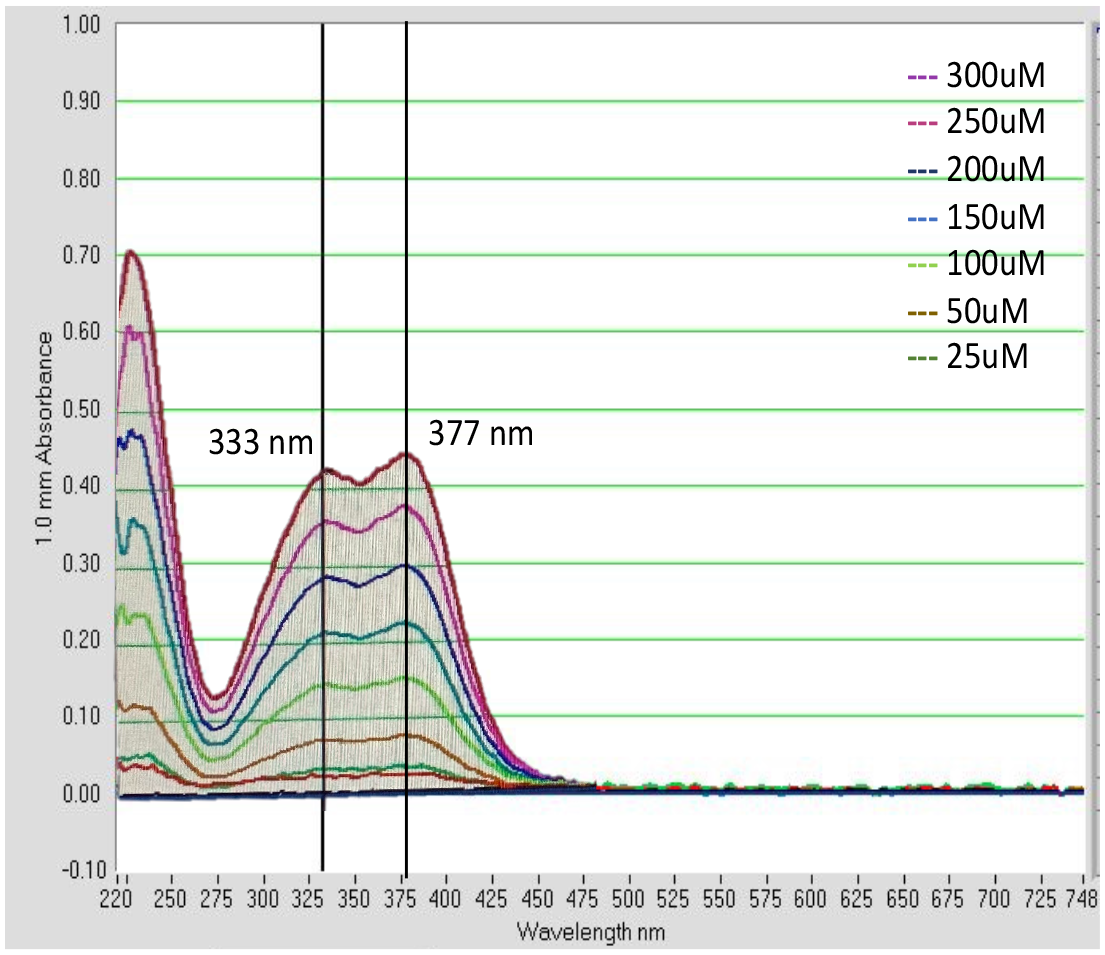
Series of UV/VIS spectra at pH 9.3 for each niclosamide solution corresponding to the photographic images of the vials in ***Figure S6***. The absorbance of the peak at 333nm was used in the calibration (although the maximum in aqueous pH 9.3 buffer was closer to 335nm).

***Figure S8*** shows the UV/Vis Absorption calibration at 333nm (averaged for at least 5 measurements) and plotted versus Niclosamide Solution Concentration (uM). With a blank measured at a maximum of 0.003 absorption units, the line is given by y = 0.0014x + 0.0006. Each absorbance value (Abs_Nic_) was therefore converted to niclosamide concentrations [Nic] (uM) using this calibration equation, i.e., [Nic] = (Abs_Nic_ −0.0006)/0.0014. Equivalent values for ug/mL are given in the table inset.

**Figure S8.**
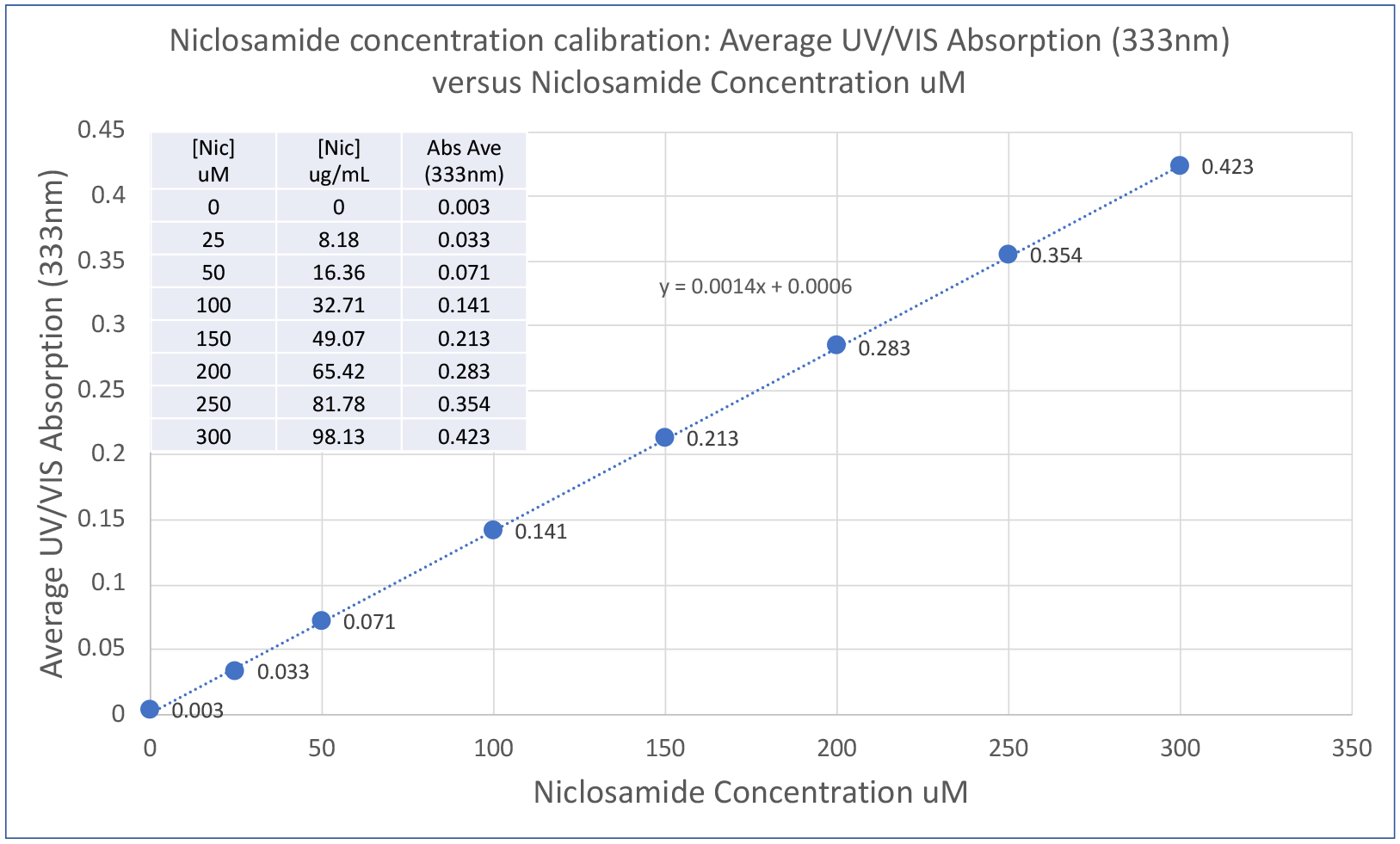
UV/VIS Calibration for niclosamide solutions. Averaged UV/VIS Absorbance at 333nm for Niclosamide Solutions. Data labels give the absorbance at each niclosamide concentration.

## Notes

### Competing Interest Statement

The authors have declared no competing interest.

### Summary of Updates

Updates in response to a review critique including some clarifications to terms. Also in materials, more details are given for AKSci niclosamide purity and stability. A slight title change reflects the potential universality of the niclosamide solution

